# Cell Type Specific CAG Repeat Expansions and Toxicity of Mutant Huntingtin in Human Striatum and Cerebellum

**DOI:** 10.1101/2023.04.24.538082

**Authors:** Kert Mätlik, Matthew Baffuto, Laura Kus, Amit Laxmikant Deshmukh, David A. Davis, Matthew R. Paul, Thomas S. Carroll, Marie-Christine Caron, Jean-Yves Masson, Christopher E. Pearson, Nathaniel Heintz

## Abstract

Brain region-specific degeneration and somatic expansions of the mutant Huntingtin (mHTT) CAG tract are key features of Huntington’s disease (HD). However, the relationships between CAG expansions, death of specific cell types, and molecular events associated with these processes are not established. Here we employed fluorescence-activated nuclear sorting (FANS) and deep molecular profiling to gain insight into the properties of cell types of the human striatum and cerebellum in HD and control donors. CAG expansions arise in striatal medium spiny neurons (MSNs) and cholinergic interneurons, in cerebellar Purkinje neurons, and at *mATXN3* in MSNs from SCA3 donors. CAG expansions in MSNs are associated with higher levels of MSH2 and MSH3 (forming MutSβ), which can inhibit nucleolytic excision of CAG slip-outs by FAN1 in a concentration-dependent manner. Our data indicate that ongoing CAG expansions are not sufficient for cell death, and identify transcriptional changes associated with somatic CAG expansions and striatal toxicity.

## INTRODUCTION

Huntington’s disease (HD) is a fatal neurodegenerative disease characterized by delayed age at onset, vulnerability of specific neuronal populations to degeneration, and accumulation of protein aggregates, features that are shared with other, more prevalent late-onset neurodegenerative diseases. HD is caused by an abnormally long CAG tract in exon 1 of the Huntingtin gene (HTT). The mutation is autosomal dominant, variably penetrant if the CAG tract is 36 - 39 repeat units long, and fully penetrant at repeat lengths of 40 CAGs and above^1^. Abnormalities in affected patients include motor, cognitive and psychiatric problems that arise earlier or manifest at higher frequency in patients with longer CAG tracts^2, 3^. HD age at onset is most often defined as the onset of motor symptoms, which are thought to arise as a consequence of early degeneration of the caudate nucleus and putamen, primarily due to loss of projection neurons of these structures, known as medium-sized spiny neurons of the direct and indirect pathways (dMSNs and iMSNs)^4^. Remarkably, other neuron types within these same structures are largely spared from cell death^5–7^. The mutant *HTT* allele (*mHTT*) encoded CAG repeats give rise to a long polyglutamine stretch that makes the mHTT protein prone to the formation of cytoplasmic and nuclear aggregates^8^. However, *HTT* is ubiquitously expressed and the reason for selective vulnerability of specific cell types in HD is largely unknown^9^.

The *mHTT* CAG repeat is unstable intergenerationally and somatically^10, 11^. Expansion of the inherited *mHTT* allele to very long CAG tracts has been observed sporadically in various brain structures, including the caudate nucleus and putamen^12^. A causal role for somatic expansions of the CAG repeat in HD pathogenesis is supported by two findings from a genome-wide association study looking for genetic modifiers of HD motor symptom onset other than CAG tract length itself^3, 13^. First, CAG repeat length predicts the age at motor symptom onset better than polyglutamine tract length. Second, several of the candidate genes identified in these studies are known to be involved in DNA repair. The latter finding is consistent with previous studies demonstrating that several of these DNA repair genes alter somatic expansions of long CAG repeats in brain tissue of transgenic mouse models of HD^14–16^. These data have led to a two-component model of HD pathogenesis that first involves somatic expansion of the *mHTT* CAG tract, followed by a toxic effect of the mHTT protein encoding an expanded polyglutamine tract^17^. Although this model is consistent with most studies of HD mouse models and human neuropathology, it is not known why somatic CAG expansions occur in specific brain regions, whether CAG expansions occur in specific cell types in these regions, whether CAG expansions are sufficient to explain selective cellular vulnerability in HD, and if cell-specific factors in addition to somatically expanded *mHTT* CAG tract are required for toxicity.

To gain further insight into somatic CAG expansion and toxicity in HD, we have developed FANS methods for isolation of large numbers of nuclei from human striatal and cerebellar cell types, and examined the relationships between selective cellular vulnerability, somatic CAG expansion, and transcriptional responses in HD. We find that extensive somatic expansion of the *mHTT* CAG tract occurs in both MSN populations that are selectively vulnerable in HD, as well as in cholinergic interneurons that are not lost in the HD striatum. CAG expansion is observed also at the mutant *ATXN3* locus in MSN nuclei isolated from the post-mortem brains of spinocerebellar ataxia 3 (SCA3) donors, suggesting that MSNs are intrinsically prone to somatic expansion of CAG tracts. We demonstrate that the levels of DNA mismatch repair proteins MSH2 and MSH3 are elevated in MSN nuclei, suggesting that these proteins may contribute to preferential somatic CAG expansions in MSNs. We offer mechanistic insight to how MutSβ could be promoting somatic CAG expansions by showing that increased concentrations of MutSβ inhibit excision rates of excess slipped-CAG repeats, intermediates of expansion mutations, by FAN1 nuclease. The *mHTT* CAG tract also undergoes selective expansion in Purkinje cells relative to other cerebellar cell types, consistent with the reports of Purkinje cell loss in HD and other CAG repeat expansion disorders. Our data demonstrate that somatic CAG expansion and mHTT toxicity are not inextricably coupled in human striatal cell types, and they identify transcriptional programs that are associated with selective cellular vulnerability and toxicity in the human HD brain.

## RESULTS

### FANSseq profiling of human striatal cell types

To characterize the expression profile, somatic mutations and chromatin accessibility of human striatal cell types, we further developed the method of fluorescence-activated nuclei sorting (FANS)^18^ for isolating nuclei from human post-mortem caudate nucleus and putamen (**Figure 1A**). Nuclei were purified from samples dissected from human post-mortem caudate nucleus and putamen, stained with either antibody or RNA probes specific for the nuclei of cell types of interest, and resolved by passage through a FACS (**Figure 1A**) (**Table S1**). We collected nuclei of oligodendrocytes, astrocytes, microglia, dopamine receptor D1-expressing (DRD1+) medium spiny projection neurons of the direct pathway (dMSNs), dopamine receptor D2-expressing (DRD2+) medium spiny projection neurons of the indirect pathway (iMSNs) and 4 different types of striatal interneurons: cholinergic interneurons expressing choline acetyltransferase (CHAT+ INs), fast-spiking interneurons expressing Parvalbumin (PVALB+ INs), somatostatin-expressing interneurons (SST+ INs) and primate-specific Tachykinin Precursor 3-expressing interneurons (TAC3+ INs)^19^ (**Figures 1B and S1**). The specificity of labeling probes and purity of isolated populations of nuclei were verified by generating RNA-seq libraries from their nuclear transcriptomes (FANSseq) and comparing their gene expression profile to well-known cell type markers and previously published data^19, 20^ (**Figures 1C and 2A**).

**Figure 1.**
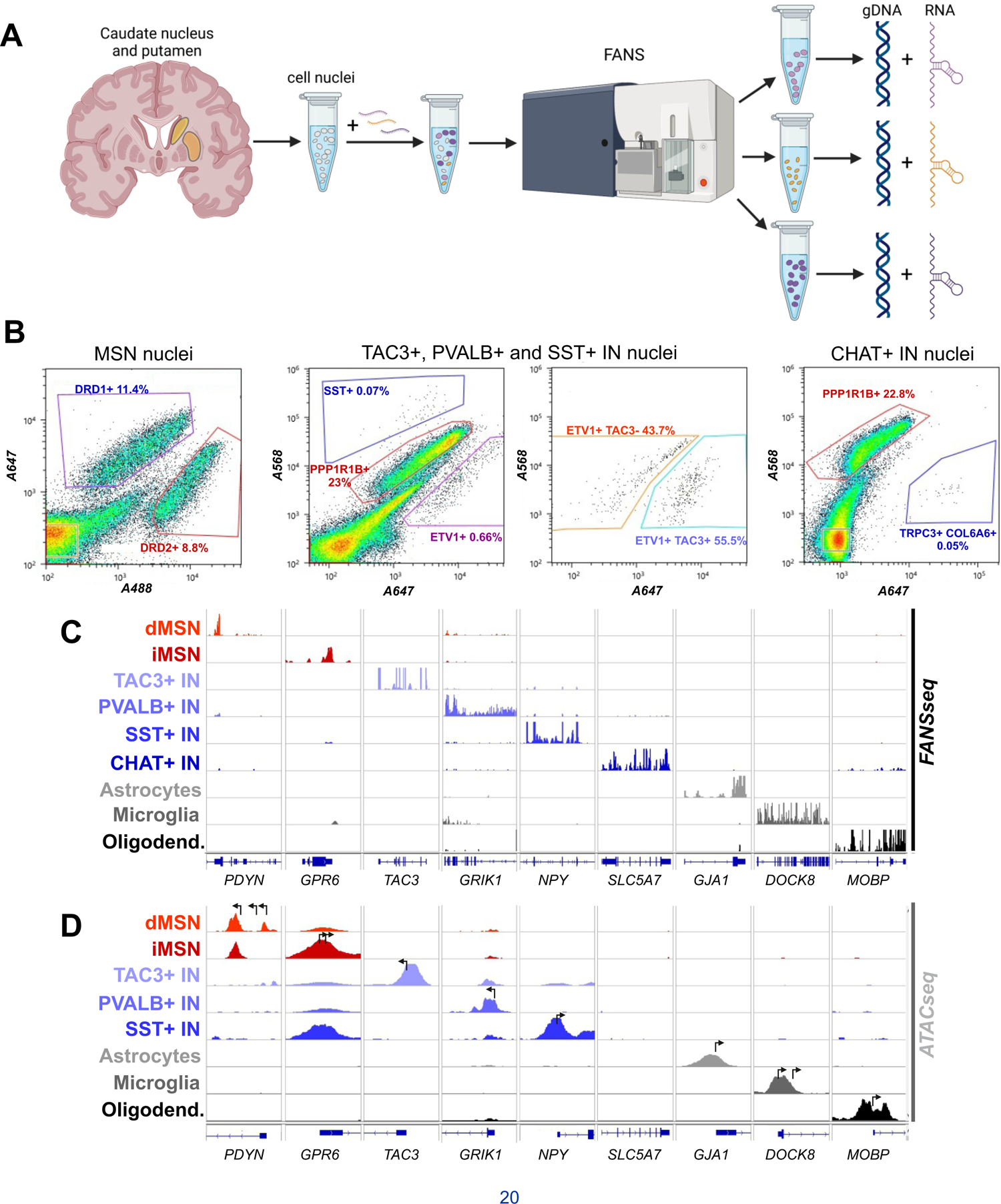
FANS-based isolation of nuclei of striatal cell types from human post-mortem caudate nucleus and putamen. (**A**) Schematic representation of the procedure used to extract cell type-specific genomic DNA and nuclear RNA from cell nuclei labeled with cell type-specific probes. Created with BioRender.com. (**B**) Representative FANS plots showing the labeling of striatal cell nuclei with PrimeFlow probes specific for dMSNs (DRD1+ nuclei), iMSNs (DRD2+ nuclei), SST+ interneurons (SST+ nuclei), PVALB+ interneurons (ETV1+ TAC3-nuclei), TAC3+ interneurons (ETV1+ TAC3+ nuclei) and CHAT+ interneurons (TRPC3+ COL6A6+ nuclei). The probe against *PPP1R1B* labels all MSN nuclei. The detailed strategy used for sorting is described in Materials and Methods. (**C-D**) Representative distribution of human FANSseq (**C**) and ATACseq (**D**) reads mapped to genes expressed specifically in each of the striatal cell types studied. In **D**, arrows mark the position of annotated transcriptional start sites. The data are from a 41-year-old male control donor.

**Figure 2.**
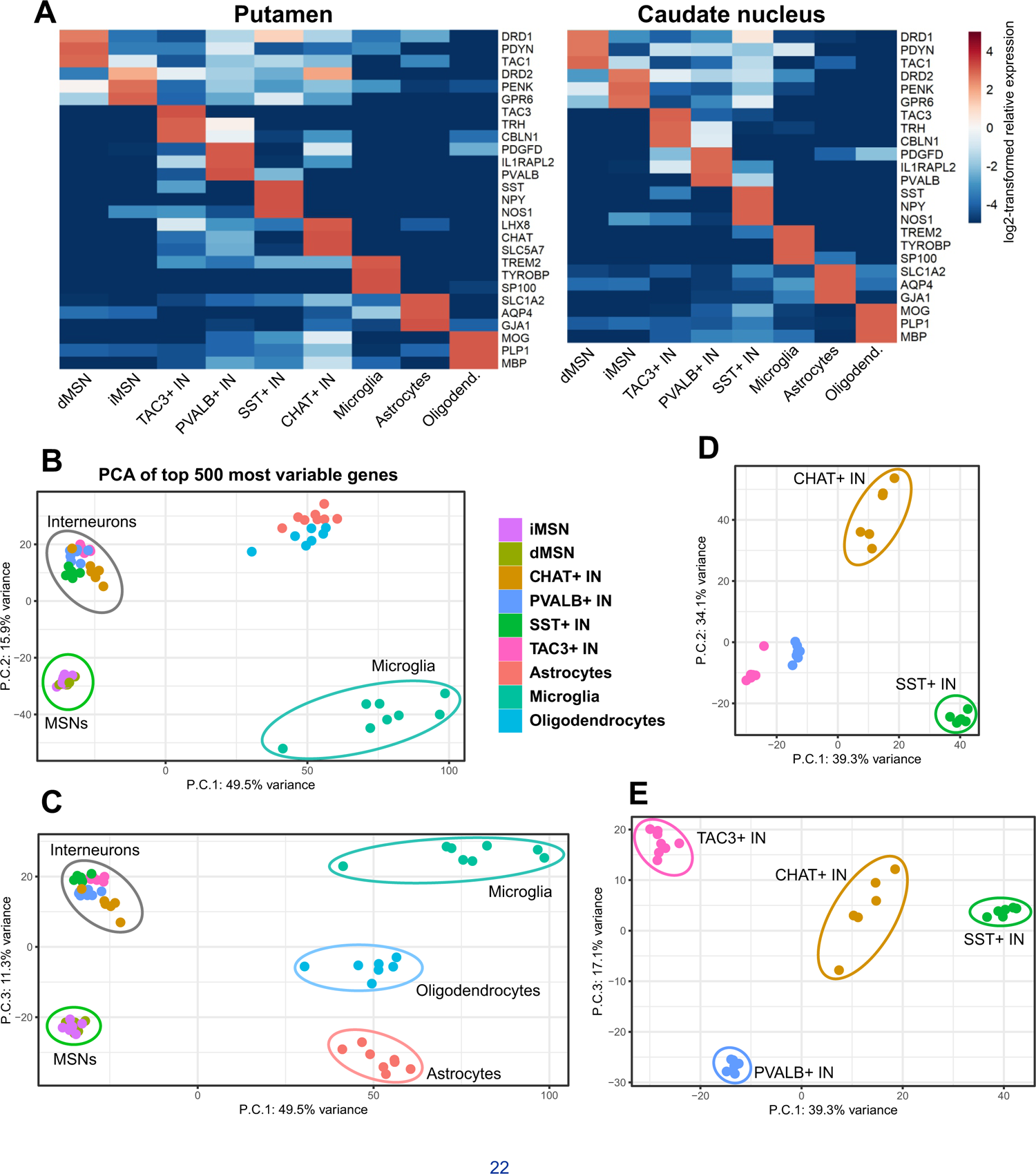
Purity and reproducibility of the isolation of striatal cell type nuclei across the two striatal brain regions studied. (**A**) Heatmaps depict log2-transformed relative expression level of cell type-specific marker genes in each cell type, calculated based on the mean of DESeq2-normalized counts from 6-8 control donors. (**B-E**) Principal component analysis of FANSseq data from all putamen cell types (**B and C**) or interneuron types (**D and E**) from 6-8 control donors.

Analysis of main sources of variance between FANSseq datasets from all putamen cell types indicated that the first three principal components (P.C.) separated neuronal datasets from glial ones (P.C. 1), MSN datasets from those of interneurons (P.C. 2) and datasets of different glial cell types from each other (P.C. 3) (**Figures 2B and 2C**). For FANSseq datasets from putamen interneurons the major principal components separated the datasets according to interneuron subtype (**Figures 2D and 2E**). We asked how the interneuron types collected by our FANS-based methodology corresponded to the human striatal interneuron types recently defined via single-nucleus RNA-seq (snRNA-seq)^19^. A majority of marker genes with enriched expression in a specific interneuron type in snRNA-seq data had also enriched expression in our corresponding FANS-based interneuron populations (**Figure S2**). Analysis of variance between FANSseq datasets from MSNs of both the caudate nucleus and putamen indicated that the main principal components separated datasets according to MSN subtype (P.C.1) and donor sex (P.C.2) (**Figure S3A**). Notably, none of the top principal components related to the brain region of origin (i.e. caudate nucleus or putamen) (**Figure S3A**).

Important features of the FANS-based molecular studies of post-mortem human tissue are the ability to isolate tens of thousands of nuclei from the same cell type, to generate deep transcriptional profiles, and the high sensitivity that can be achieved in epigenetic studies of chromatin accessibility ^18^. Given the recent demonstration that ambient RNA released from the cytoplasm or nucleus is a source of contamination in single nucleus RNA-seq studies of brain tissue^21^, the difficulty in setting an arbitrary TPM (transcripts per million) threshold for gene expression in RNAseq datasets, and the likelihood that low-level ambient RNA can confound studies of differential gene expression, we used promoter accessibility in ATACseq data as an independent measure of active genes^22^. To this end, we excluded genes for which none of their annotated transcriptional start sites overlapped with regions of accessible chromatin in that cell type, as defined by the presence of a consensus peak in ATACseq data generated from these cells (**Figure 1D**). We reasoned that this would allow us to minimize the number of possible false-positive differences in gene expression that might have resulted from contaminating ambient transcripts, FACS sorting impurities or contamination with genomic DNA. We were not able to produce ATAC-seq datasets from CHAT+ INs due to their low abundance in striatal tissue. Based on the criterion of promoter accessibility, the average number of genes expressed in each striatal cell type is about 13,000 – 14,000 (**Figure S3B**). This estimate roughly corresponds to the average number of genes with expression level above an arbitrary cutoff value of 5 transcripts per million (TPM) in individual FANSseq datasets of neuronal cell types (**Figure S3C**). Therefore, the individual datasets capture most of the transcriptome complexity of these cell types at a well-measurable level. In summary, we have used coordinated FANSseq and ATACseq profiling to generate comprehensive high-quality datasets for each neural cell type present in the human caudate nucleus and putamen.

### Somatic expansions of the *mHTT* CAG repeat occur in vulnerable MSNs and resilient CHAT+ INs

Tissue-specific ongoing CAG repeat expansions of the mutant allele are a central feature of HD and other repeat expansion disorders^11, 23–26^. Large somatic expansions of the *mHTT* exon 1 CAG tract in the striatum and cerebral cortex have been demonstrated by small pool PCR^12^, and laser capture studies have suggested that somatic expansion can occur in both neurons and glia, albeit more frequently in neurons^27^. To address the specificity of somatic expansion of *mHTT* CAG tract quantitatively, and to understand whether it is correlated with cell loss in HD, we isolated nuclei of each striatal cell type by FANS (**Figure S4**), verified the purity of isolated populations of nuclei by analysis of marker gene expression in nuclear transcriptomes from the isolated nuclei (**Figures S5A and S5B**), and measured the length of *HTT* exon 1 CAG tract in the isolated genomic DNA from these populations using Illumina-sequencing of amplicons derived from *HTT* exon 1^28^. For the great majority of samples, genomic DNA from 800 or more cells was used for CAG tract length measurement.

Analysis of gDNA of different cell types from five HD donors carrying most prevalent disease-causing CAG tract lengths (from 42 to 45 uninterrupted CAGs, see **Figure S5A**) revealed that *mHTT* CAG tract was relatively stable in glial cell types and SST+, TAC3+ and PVALB+ INs (**Figures 3A-3C**). However, only a small fraction of dMSNs and iMSNs had *mHTT* copies with the original inherited CAG tract length and approximately half of these neurons had CAG tracts that were expanded by more than 20 repeat units (**Figures 3A-3C**). To reveal differences in the “non-expanding” striatal cell types, we calculated their ratios of somatic expansion - a metric that has been used for describing the distribution of CAG tract sizes in cells undergoing modest somatic repeat expansion^28^. These data demonstrated that striatal oligodendrocytes had consistently greater CAG expansions compared to microglia, astrocytes, and SST+, PVALB+ and TAC3+ INs (**Figure S5C**). There was no evidence of somatic expansion of the normal *HTT* allele CAG repeat tract in any of the cell types analyzed (**Figure S5D**).

**Figure 3.**
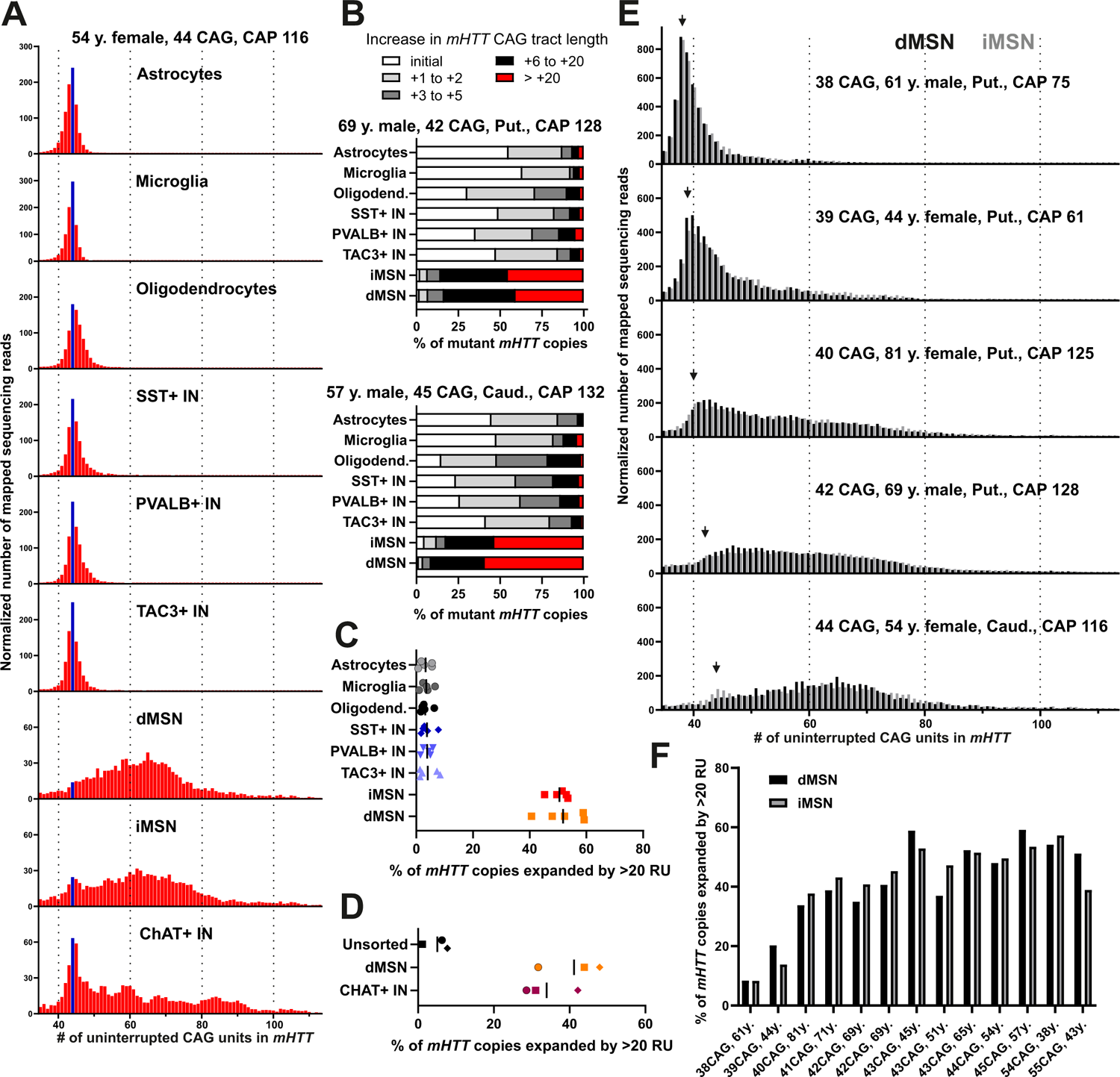
*mHTT* CAG tract undergoes somatic expansion in selected striatal neuron types. (**A**) Length distribution of *mHTT* CAG tract in studied cell types of caudate nucleus and putamen of a 54-year-old female donor that carried a tract of 44 uninterrupted CAG units. Blue bar marks sequencing reads derived from the initial unexpanded CAG tract. Y-axes denote normalized number of sequencing reads mapped to reference sequences with different CAG tract lengths (normalized by scaling to 1000 reads). Reads derived from the normal HTT allele are not shown. (**B**) Frequency distribution of *mHTT* copies with different CAG tract length increases. Data is shown for striatal cell types of two donors that carried the most common HD-causing CAG tract lengths. (**C-D**) Comparison of the fraction of *mHTT* copies with CAG tracts expanded by more than 20 repeat units (RU). (**C**) While the frequency of most highly expanded *mHTT* CAG tract copies was not different between dMSN and iMSN, comparison of each of these to any other striatal cell type showed a statistically significant difference (P < 0.0001 by one-way ANOVA, adjusted P value <0.0001 in Holm-Sidak’s multiple comparisons test). (**D**) The frequency of most highly expanded *mHTT* CAG tract copies was not different between dMSNs and CHAT+ interneurons, but comparison of each of these to unsorted nuclei showed a statistically significant difference (P = 0.0013 by one-way ANOVA, adjusted P value <0.004 in Holm-Sidak’s multiple comparisons test). Different symbols are used for each of the three donors, all carrying *mHTT* allele with an initial length of 44 uninterrupted CAG repeats. (**E**) Length distribution of *mHTT* CAG tract in MSNs of caudate nucleus and putamen in donors carrying *mHTT* alleles of reduced and full penetrance. Arrowhead indicates the initial unexpanded size of the CAG tract. (**F**) Comparison of the fraction of *mHTT* copies with CAG tracts expanded by more than 20 repeat units (RU). P = 0.896 between cell types, in ratio paired t-test.

Because the scarcity of tissue available prevented us from isolating striatal CHAT+ IN nuclei from all but one of these five initially characterized HD donors, we isolated nuclei of this very rare cell type from two other HD donors. In these samples the fraction of nuclei with large repeat length increases was large, revealing that *mHTT* CAG tract undergoes large expansions also in CHAT+ INs (**Figures 3A and 3D**).

Pairwise comparisons of the two MSN types from donors carrying a wider range of initial repeat lengths revealed that the extent of repeat expansion was dependent on initial repeat length. Thus, more modest expansion was observed for *mHTT* alleles with reduced penetrance (CAG tract lengths < 40 repeats) relative to longer, fully penetrant alleles (**Figures 3E and 3F**). Interestingly, while iMSNs have been reported to be more vulnerable of the two MSN subtypes^29^, there was no significant difference in the frequency of highly expanded repeat copies between dMSNs and iMSNs (**Figures 3E and 3F**).

Taken together, the *mHTT* CAG tract expands dramatically in striatal MSNs and CHAT+ INs, and very little in other cell types analyzed. These data support the hypothesis that extensive somatic expansion of the *mHTT* CAG tract is required for the vulnerability of MSNs in HD. However, given previous studies demonstrating that CHAT+ INs in the striatum are not lost in HD^5, 6^, it is also evident from our results that robust expansion of the *mHTT* CAG tract is not sufficient to cause neuronal loss in HD (see also the accompanying manuscript by Pressl et al). In addition, our data show that differences in the rate of *mHTT* CAG expansion are an unlikely explanation for the reportedly greater vulnerability of iMSNs than dMSNs in this disease.

### Cell type-specific instability of the *mHTT* CAG tract in the HD cerebellum

The loss of cerebellar Purkinje cells (PCs) in several spinocerebellar ataxias (SCA1, SCA2, SCA6 and SCA7) where the causal elongated CAG tracts undergo germline expansion has suggested that in these disorders, somatic CAG expansion may occur in PCs^30^. Although the viability of PCs in HD has been a matter of debate, ataxia is not an uncommon symptom in HD patients^31^ and recent stereological studies have demonstrated that PC loss occurs in the cerebellum in HD cases with predominant motor symptoms^32^. Given these data, we measured *mHTT* CAG instability in cerebellar cell types in several HD donors (**Table S1**). To isolate the nuclei of granule cells, astrocytes, microglia, oligodendrocytes and their progenitors (OPCs), we followed the method of Xu et al. using antibodies to nuclear proteins^18^. Nuclei of the very rare Purkinje cells (<0.01% of all nuclei) were labeled with PrimeFlow probes that bind to the transcripts of carbonic anhydrase 8 (*CA8*) gene. Although the degree of *mHTT* CAG expansion in PCs was relatively modest compared to MSNs, the tract had expanded more in PCs than in any other cerebellar cell type (**Figures 4A-4C**). The variability in somatic CAG expansion observed for PC samples from different donors can most likely be attributed to the rarity of this cell type, which made it extremely difficult to collect samples entirely free of contamination with nuclei of ‘non-expanding’ cell types. Similar to striatal oligodendrocytes, mature cerebellar oligodendrocytes had more expanded CAG tracts compared to granule neurons, other glia types or OPCs within each donor, even though statistical significance was not reached for all of these comparisons (**Figure 4C**). While the degree of somatic *mHTT* CAG expansion differs between Purkinje cells and MSNs, our data indicate that both in striatum and cerebellum the *mHTT* CAG tract is somatically unstable in selected neuron types and much more stable in other neuron types and glial cells.

**Figure 4.**
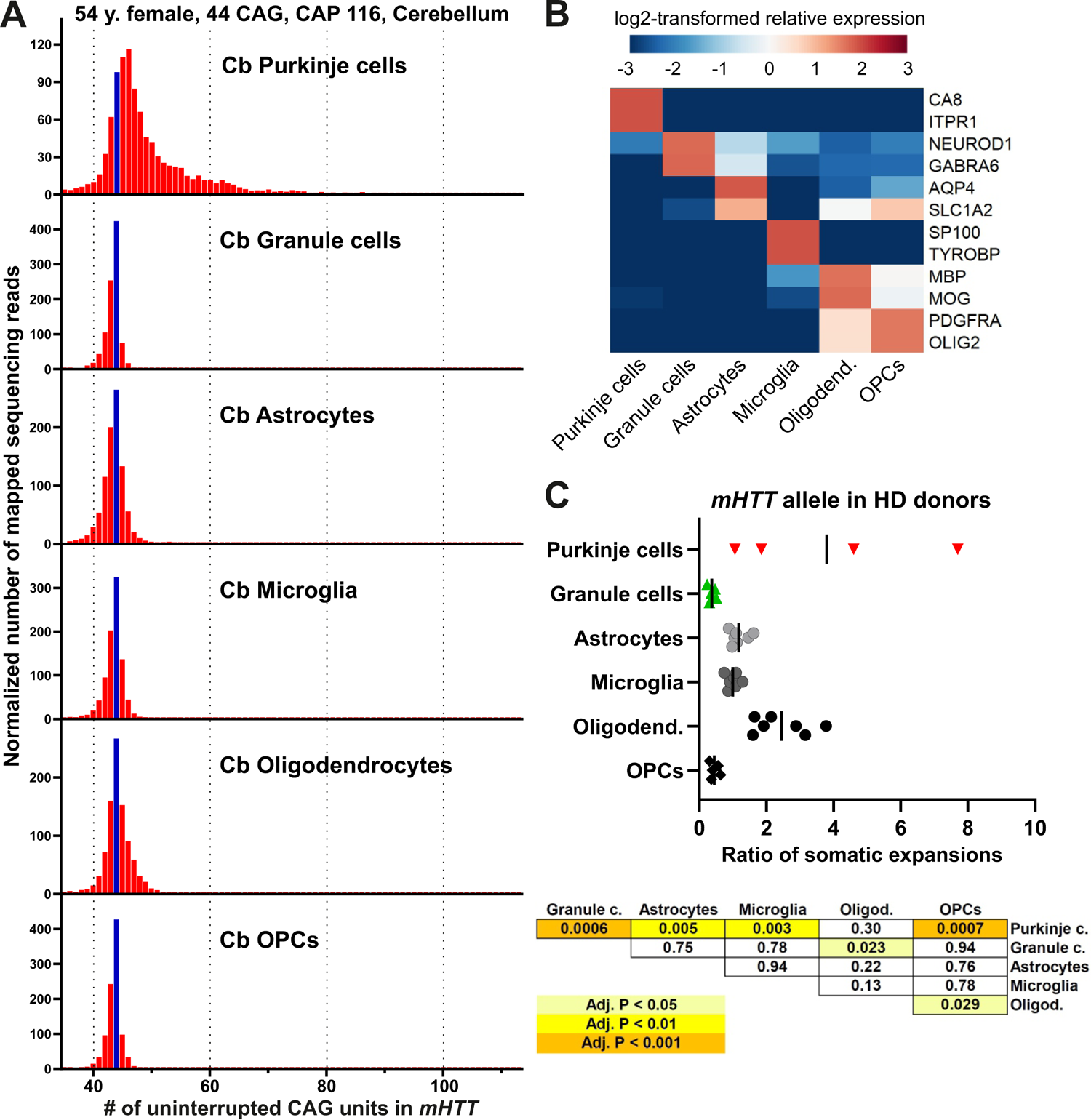
Expansion of *mHTT* CAG tract in cerebellar Purkinje cells. (**A-B**) Length distribution of *mHTT* CAG tract (**A**) and cell type marker gene expression (**B**) in cerebellar cell nuclei isolated from a 54-year-old female donor that carried a tract of 44 uninterrupted CAG units. Blue bar marks sequencing reads derived from the initial, unexpanded CAG tract. Reads derived from the shorter normal HTT allele are not shown. (**B**) Heatmap depicts log2-transformed relative expression in each sample (calculated based on DESeq2-normalized counts). (**C**) Comparison of the calculated ratio of somatic expansions of *mHTT* CAG tract in cerebellar cell types. The table presents adjusted P values as calculated by Holm-Sidak’s multiple comparisons test post one-way ANOVA (P < 0.0001).

### Striatal MSNs are prone to somatic CAG expansion

Preferential somatic expansion of the *mHTT* CAG tract in selected striatal neuron types could be due to cell type-specific properties of the *HTT* locus (i.e. MSN and CHAT+ IN-specific factors acting in *cis*), putative cell type-specific factors acting in *trans* (e.g. DNA repair proteins) or a combination of the two. Since the expansion-promoting effect of putative *trans*-factors would not necessarily be limited to the *mHTT* locus, we asked whether MSNs have a propensity to expand long and pure CAG or CTG tracts at other genomic loci as well. Because transcription through the repeat seems to be a prerequisite for somatic expansions^33^, we chose to analyze the CAG repeat in *ATXN3* gene because, as is the case for HTT, its transcription is relatively uniform across striatal cell types (**Figure S6A**).

We isolated glial cell and MSN nuclei from striatal tissue of five donors with spinocerebellar ataxia 3 (SCA3) and striatal interneuron nuclei from two SCA3 donors, all carrying a long CAG repeat in the mutant ATXN3 allele (*mATXN3*) (**Table S1**). Despite the presence of a long uninterrupted CAG repeat, with initial lengths ranging from 64 to 76 CAG repeats in the SCA3 donors analyzed, there were no clear signs of MSN loss even in the oldest SCA3 donors, as judged by the abundance of large NeuN+ nuclei in striatal homogenates (**Figure S6B**). The m*ATXN3* gene CAG tract was clearly more unstable in the MSNs relative to glial cells and interneurons (**Figures 5A and 5B**). However, despite the increased length of the inherited *mATXN3* repeat tract, its expansion was modest compared to that of m*HTT* CAG repeat tract in MSNs in HD. For example, there were very few copies of the *mATXN3* allele with repeat expanded by more than 20 repeat units (**Figure 5A**). These data indicate that MSNs have a propensity to expand long CAG tracts at other genomic loci and suggest that in addition to the CAG repeat tract length there are other locus-specific properties that are important for the dramatic expansion of the *mHTT* CAG tract. These observations support the hypothesis that *mHTT* exon 1 CAG tract instability is modulated by rate-limiting *trans*-acting factors expressed at different levels in striatal cell types.

**Figure 5:**
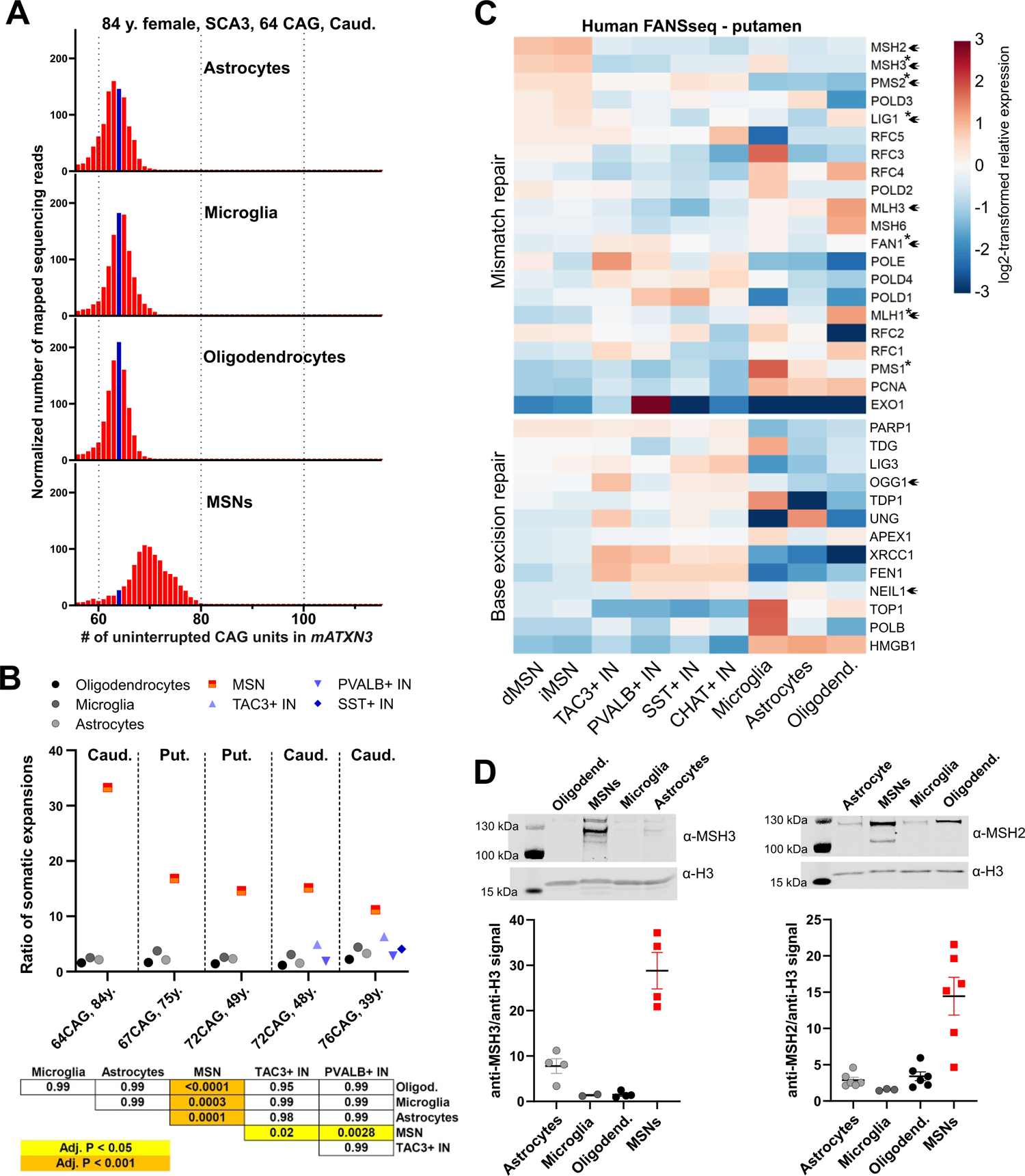
MSNs are prone to somatic expansion of *mATXN3* CAG tract and have elevated expression of nuclear MSH2 and MSH3 proteins. (**A**) Length distribution of *mATXN3* CAG tract in cell types of caudate nucleus of an 84-year-old female donor that carried a tract of 64 uninterrupted CAG units. Size of the initial unexpanded CAG tract is marked with a blue bar. Reads derived from the normal *ATXN3* allele are not shown. (**B**) Comparison of the calculated ratio of somatic expansions of *mATXN3* CAG tract in striatal cell types. The table presents adjusted P values as calculated by Holm-Sidak’s multiple comparisons test following one-way ANOVA (P < 0.0001). (**C**) Heatmaps depict log2-transformed relative expression of MMR and BER genes in cell types of putamen, calculated based on DESeq2-normalized counts from 6-8 control donors. Genes identified as HD age at onset-modifying candidates or known to influence CAG tract instability in HD mouse models are marked with an asterisk or arrowhead, respectively. (**D**) Representative immunoblots and quantification of anti-MSH3/anti-H3 (left) and anti-MSH2/anti-H3 signal ratio (right) in unfixed nuclei isolated from the putamen of 4 or 6 control donors, respectively. These ratios were higher for MSNs compared to other cell types analyzed (P < 0.0001 by one-way ANOVA, adjusted P value ≤0.0005 in Tukey’s multiple comparisons test). Data are presented as mean +/- SEM.

To identify these *trans*-acting factors that may explain preferential CAG expansion in MSNs, we compared the FANSseq expression profiles of striatal cell types. We focused on genes coding for DNA mismatch repair (MMR) and base-excision repair (BER) proteins as several of these proteins have been shown to affect repeat instability in model systems^34^, and because several MMR genes are represented among candidate genes identified as age of motor symptom onset modifiers in HD mutation carriers^13, 35^. We found that transcript levels of *MSH2* and *MSH3*, encoding MMR proteins that form the MutSβ complex, were more than 2-fold higher in both dMSNs and iMSNs compared to other striatal neurons, including CHAT+ INs, and this difference was consistent across neuron types in both putamen and caudate nucleus (**Figure 5C, and Figures S7A-S7C**).

To determine whether the FANSseq data accurately reflected nuclear protein levels for these factors in abundant striatal cell types, we measured MSH2 and MSH3 levels by Western blotting of nuclear lysates from MSNs, microglia, astrocytes and oligodendrocytes (**Figure 5D**). The ratio of both MSH2 and MSH3 to chromatin, assessed by anti-H3 signal, was significantly higher in MSN nuclei compared to glial cells (**Figure 5D**). Knockouts of *Msh2* or *Msh3* is sufficient to eliminate somatic CAG expansions in HD mouse models carrying expanded CAG tracts of over 100 repeat units, and mice heterozygous for *Msh2*+/- or *Msh3*+/- show intermediate levels of CAG expansions compared to the high levels of expansions seen with wildtype levels of these MMR proteins^15, 16, 36–38^. Moreover, naturally occurring murine genetic variants of *Msh3* that up- or down-regulate the expression of MSH3, lead to higher or lower levels of somatic CAG expansions in HD mouse models^38^. These observations suggest that elevated levels of the two components of MutSβ may explain the enhanced CAG expansions in human MSNs.

### High MutSβ levels suppress FAN1’s excision of excess slipped-CAG DNA

*FAN1* was identified as a modifier of HD disease^3^ and the nuclease activity of FAN1 has been shown to suppress CAG expansions in the central nervous system of HD mice and in cells derived from HD patients^39–44^. Thus, FAN1 is a suppressor of expansions while MutSβ is a driver of expansions. Unlike the levels of *MSH2* and *MSH3* transcripts, *FAN1* transcript levels are not higher in MSNs compared to other striatal neuron types (**Figure S7D**). We asked whether a higher MutSβ to FAN1 ratio, as predicted based on elevated *MSH2* and *MSH3* expression in MSNs, might affect the rates of excess slip-out excision by the FAN1 nuclease. To answer this, we used purified recombinant human proteins (**Figure S8A**) and slipped-(CAG)20 DNA substrates, previously demonstrated to be cleaved by both *endo*- and 5’→3’ *exo*-nucleolytic activities of FAN1 (detailed in **Figure S8B**)^44^, which was labeled on the 5’ or 3’ end to distinguish between the two activities of FAN1.

We first established conditions where FAN1 endo-nucleolytic digestion of a CAG slip-out DNA is only partially complete. Cleavage occurred just past the elbow “E” in the duplex and proximal to the 5’ end of the slip-out, with digestion products electrophoretically resolving at the top and bottom of the gel, respectively (**Figure 6A**, compare lane 1 with 2). Addition of increasing concentrations of MutSβ lead to progressive and significant inhibition of excision by FAN1 (**Figure 6A**, compare lane 2 with lanes 6-8). In contrast, addition of increasing concentrations of MutSα, a dimer of MSH2 and MSH6, did not inhibit cleavage significantly (**Figure 6A**, compare lane 2 with lanes 3-5). This result agrees with the observation that MSH6 is not involved in somatic CAG expansions^15^. The limited amounts of stimulated nuclease digestion at low levels of MutSα and MutSβ are likely the result of molecular crowding, as both MutSa and MutSb preparations are devoid of nuclease activity (**Figure 6A** compare lanes 4 and 6 with lanes 9 and 10). Slipped-(CTG)20 DNA substrate could also be excised by FAN1, and this was inhibited significantly by MutSβ, but not MutSα (**Figure S8C, panel i**). Next, we tested the effect of the MutS complexes on FAN1s exo-nucleolytic digestion of slip-out DNA substrates, where “nibbling-like” cleavage occurred through-out the repeat tract (**Figures 6B and S8C panel ii**, compare lane 1 with lane 2). Exo-nucleolytic cleavage of both slipped-CAG(20) and slipped-(CTG)20 was inhibited by MutSβ, but not MutSα (**Figures 6B and S8C panel ii**, compare lane 2 with lanes 6-8 and lanes 3-5). Thus, unlike MutSα, MutSβ inhibits FAN1’s *exo-* and *endo*-nucleolytic excision of excess CAG and CTG slip-outs.

**Figure 6:**
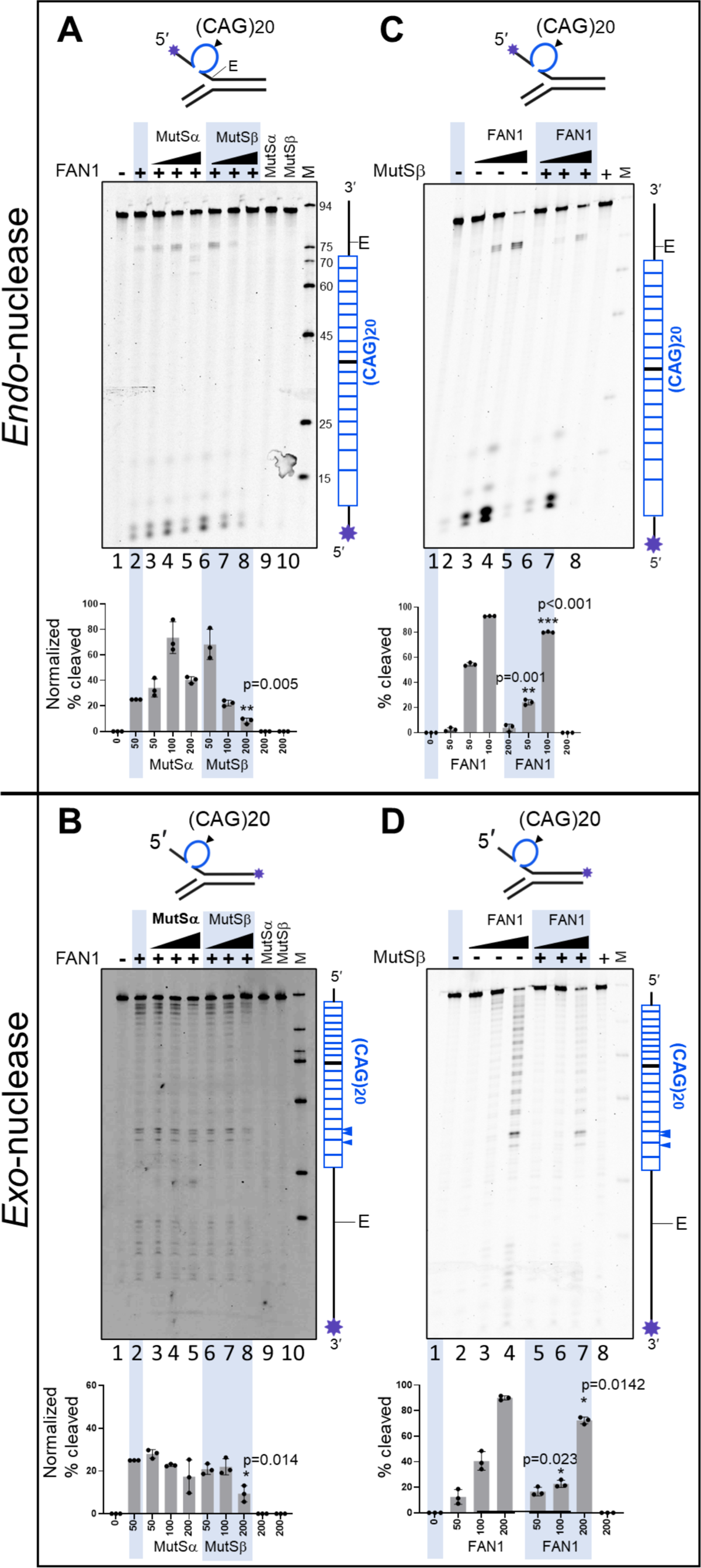
MutSβ and FAN1 levels competitively affect FAN1’s nuclease excision of excess slipped-CAG DNAs. (CAG)20-slip-out DNA substrates (schematics) mimic intermediates of expansion mutations^44^. *Endo*- and *exo*-nucleolytic activities can be distinguished by fluorescein amidite (FAM)-labeling at 5’ or 3’ ends of the (CAG)20 strand, respectively (indicated by an asterisk); in this manner, only the labeled strand and it’s digestion products are tracked. “E” is elbow at the dsDNA-ssDNA junction. (**A-B**) MutSβ not MutSα inhibits FAN1. Protein-free undigested slip-out DNA substrate (100 nM), lane 1. Slip-outs were pre-incubated with buffer, lane 2, or increasing concentrations of purified human MutSα (50, 100 or 200 nM), lanes 3-5; or MutSβ (50, 100 or 200 nM), lanes 6-8. Nuclease digestions were initiated by addition purified human FAN1 (50 nM). Lanes 9 & 10 have slip-out DNA and only MutSα (50 nM) or only MutSβ (50 nM), ensuring these purified proteins are nuclease-free. (**C-D**) Increasing FAN1 concentration can overcome MutSβ-mediated inhibition of cleavage. Protein-free undigested slip-out (100 nM), lane 1. Slip-outs were pre-incubated with buffer, lanes 1-4 or with MutSβ (200 nM), lanes 5-7. Nuclease digestions were initiated by adding increasing amounts of FAN1 (50, 100 or 200 nM), lanes 2-4 and 5-7. Lane 8 has slip-out and only MutSβ (50 nM). For gels A and B, the percentage cleavage for each reaction was normalized to cleavage levels with FAN1 alone (lane 2), and these levels were graphed (GraphPad prism 9.1). For gels C and D, the percentage cleavage for each reaction were graphed. The vertical schematic to the right of each gel indicates the location of cleavage sites along the FAM-labeled DNA strand. N = 3 replicates, mean+/- SD plotted.

Next, we tested whether excess levels of FAN1 alone or in the presence of a constant level of MutSβ affect the rates of slip-out DNA excision. Increasing levels of FAN1 rapidly digested the slip-out DNA (**Figure 6C and 6D**, compare lane 1 with lanes 2-4). While the effect of added MutSβ was evident as partial inhibition of cleavage by FAN1 (**Figure 6C and 6D**, compare lane 3 to 6 and 4 to 7), addition of increasing concentrations of FAN1 led to increased *endo*- and *exo*-nucleolytic digestion of slip-out DNA even in the presence of MutSβ (**Figure 6C and 6D**, compare lane 5 with lanes 6 & 7). FAN1 activity may, over time, be inhibited by MutSβ through competitive DNA binding, but is unlikely to involve a direct interaction of the proteins, as FAN1 does not interact with MutSβ^42^. This suggests that increased concentrations of one protein may compete with the other for the DNA substrate. To test this hypothesis, we designed a competition experiment (outlined in **Figure S8D, panel i**) incubating slipped-DNA substrates, labeled for the detection endo- or exo-nucleolytic cleavage, with MutSβ and a low concentration of FAN1 under non-digesting conditions (on ice), thus allowing MutSβ and FAN1 to competitively bind the slipped-DNA. We then split the reaction into two halves, adding BSA as protein control to one half and flooding the other half with a 5-fold excess FAN1, shifting to digesting conditions (37°C) and removing aliquots for analysis at 5-, 10-, 20- and 40-minute timepoints. Reactions with excess FAN1 progressed faster and overcame the MutSβ-mediated inhibition of *endo*- and *exo*-nucleolytic activity of FAN1 (**Figure S8D ii and iii**, compare lanes 7-11 with lanes 2-6, in both panels ii and iii). Thus, the MutSβ-mediated FAN1 inhibition is rescued by excess of FAN1. Taken together our results support a model where CAG and CTG slip-out DNA excision rates are determined by competitive binding to either MutSβ or FAN1, thereby offering an explanation to how differences in the relative level of MutSβ to FAN1 could result in CAG expansion or stabilization in different cell types.

### Expression of HD age at onset modifier candidate genes in human striatum

Studies of gene expression in animal models of human disease and post-mortem human tissue samples, as well as proteomic studies of protein complexes accumulating during pathological progression, have emerged as valuable approaches for the formulation of etiological hypotheses. To gain further insight into the pathophysiological processes that influence HD progression, we next studied the expression pattern of the remaining genes that were identified as candidates potentially modulating HD age at onset^13^. As expected, we found that candidate genes involved in MMR (marked with an asterisk on **Figures 5C and S7A**), as well as those lacking an obvious association with DNA repair are expressed at significant levels in human MSNs (**Figures S9A and S9B**). Notable exceptions are *RRM2B* and *SYT9* which are expressed at very low levels in MSNs, but at higher levels in microglia (*RRM2B*) and oligodendrocytes (*SYT9*) in both caudate nucleus and putamen (**Figures S9A, S9B and S9D**). These FANSseq results are supported by considerably lower chromatin accessibility at the promoter regions of *RRM2B* and *SYT9* in human MSNs compared to glial cell types (**Figure S9E**). This result could not have been predicted from published cell type-specific gene expression analysis of mouse striatum (**Figure S9C**). Thus, although confirmation of these data will require additional studies, our results suggest that genetic variants near *RRM2B* and *SYT9* may act in glia to influence disease onset through mechanisms that do not impact somatic expansion of the *mHTT* CAG tract.

### Altered gene expression during Huntington’s disease progression

To gain further insight into the molecular events that may play a role in somatic expansion or contribute to mHTT toxicity, we sequenced the nuclear transcriptomes of striatal MSNs and TAC3+, SST+ and PVALB+ INs from the putamen or caudate nucleus of 6-7 HD donors (**Table S1**). We again limited the comparative analysis of HD and control donors’ (n=8) FANSseq data to genes that are more likely to be truly expressed in the cell type of interest by excluding transcripts for which there was no promoter accessibility as indicated by ATACseq (**Figure S10A**). More than 98 % of genes with accessible promoters in HD MSNs had accessible promoters also in MSNs from control donors. Due to the low abundance of striatal interneurons and limited amounts of striatal tissue available from HD donors, we were not able to produce ATAC-seq datasets from the interneurons of HD donors. Therefore, striatal interneuron FANSseq datasets were filtered to include only those genes which had accessible promoters in control donors.

Principal component analysis of FANSseq data from MSNs indicated that disease status (P.C.1) and MSN subtype (P.C.2) were the main sources of variance in these datasets (**Figure S10B**). The overall number of genes with statistically significant disease-associated differences in transcript levels in each cell type is shown in **Figure 7A**. Although transcriptional changes occurred in both interneurons and MSNs, the number of both up- and downregulated genes was higher in MSNs. As expected, disease-associated changes were well correlated between dMSNs and iMSNs, but the correlation was poor in comparisons involving other neuron types (**Figure S10C**). This result shows that the majority of disease-associated transcript levels changes are not common to all striatal neuron types.

**Figure 7.**
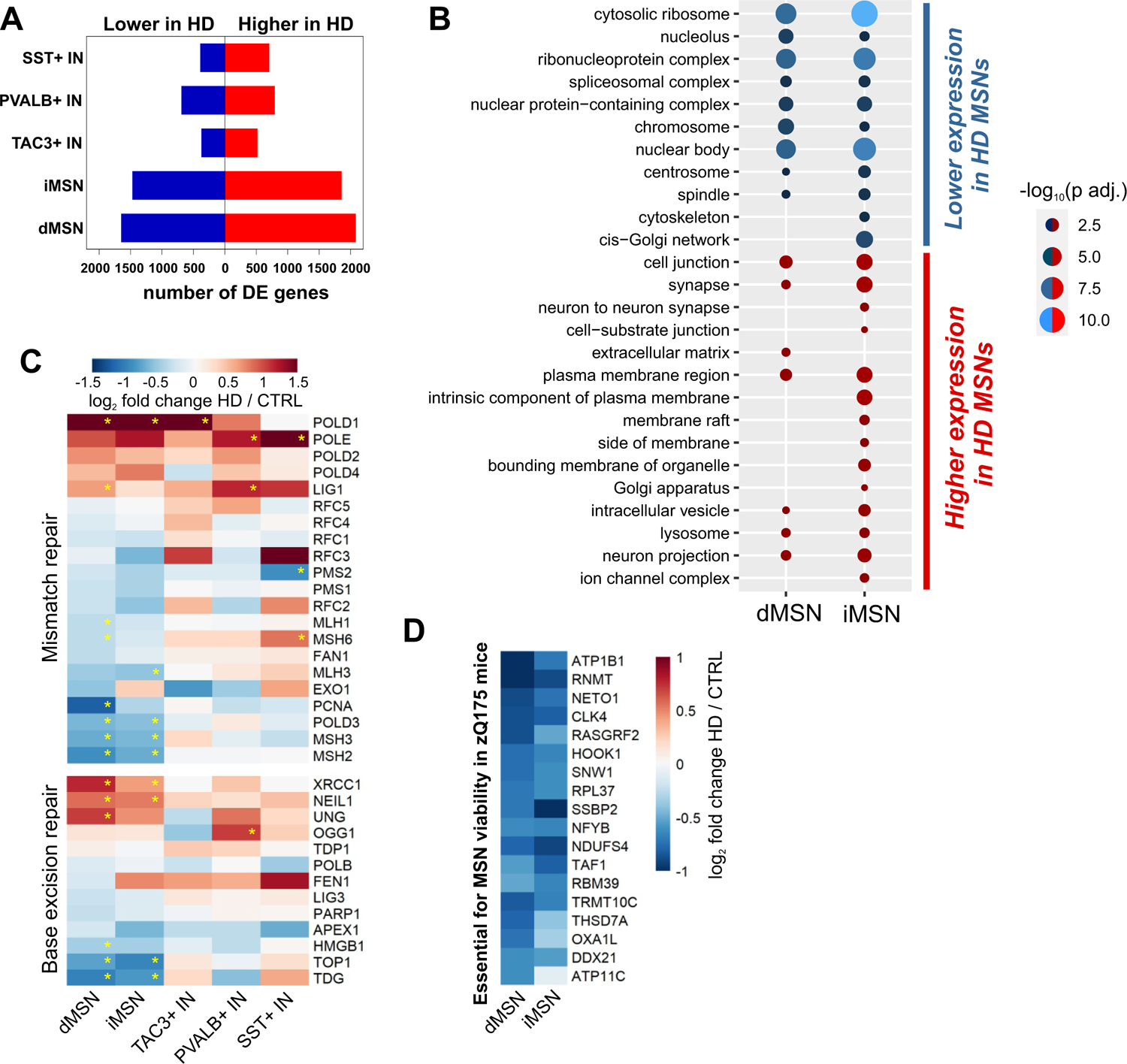
Disease-associated gene expression changes in striatal neuron types. (**A**) Number of differentially expressed (DE) genes (p adj. < 0.05 by DESeq2) in the comparison of HD and control donor FANSseq datasets from putamen or caudate nucleus. (**B**) Selected non-redundant GOCC terms from enrichment analysis of genes with disease-associated expression changes in iMSN or dMSN (p adj < 0.05), but not in any interneuron type (p adj > 0.05 in all HD vs CTRL interneuron comparisons). (**C**) Heatmaps depicting disease-associated changes in transcript levels of MMR and BER genes. Statistically significant differences are marked with an asterisk (p adj. < 0.05 by DESeq2). (**D**) Genes essential for MSN viability in zQ175 mouse model of HD that are also downregulated in HD iMSNs or dMSNs by more than a third (log2FC<-0.6, p<0.01).

### Gene ontology analysis

To identify cellular processes that are affected by disease-associated gene expression changes that take place only in MSNs, thereby correlating with the presence of more toxic mHTT species in these neurons, we analyzed which Gene Ontology Cellular Component (GOCC) terms were enriched for genes that were up- or down-regulated in MSNs but did not display these changes in expression in any of the interneuron populations studied. The results indicated that many genes downregulated specifically in MSNs are involved in ribosomal biogenesis (GOCC terms ‘cytosolic ribosome’ and ‘nucleolus’), pre-mRNA maturation (GOCC terms ‘nuclear body’ and ‘spliceosomal complex’), and other nuclear functions (**Figure 7B**). Although transcripts of mitochondrial oxidative phosphorylation pathway genes have been reported to be downregulated in HD MSNs^45^, we noticed that this disease-associated change is much more evident in the nuclear transcriptome of PVALB+ striatal interneurons (**Figures S11A and S12A**).

The GOCC terms enriched for genes that increase expression in MSNs in the HD donor data include many terms that indicate alterations in membrane protein function. ‘Neuron projection’, ‘Synapse’ and ‘Lysosome’ were among GOCC terms enriched for genes upregulated specifically in both MSN subtypes (**Figure 7B**). In contrast, for one or more of the striatal interneuron types these gene ontology categories were instead enriched for downregulated genes (**Figures S11A and S12B**). We observed also that genes central to the regulation of lysosomal biogenesis and autophagy were among the top upregulated genes in MSNs. For example, the transcripts of transcription factor TFEB, which has been shown to be essential for regulation of many genes in these pathways^46^, are strongly elevated in HD MSNs (**Figure S11B**). Several genes encoding proteins essential for autophagy are also induced in HD, including *ATG9B*, *ATG9A,* the gene encoding HTT-interacting protein ULK1 involved in autophagosome formation^47^, *MAPK8IP1* and *MAPK8IP3* which encode proteins involved in retrograde transport of autophagosomes^48^ and *SQSTM1*, the gene encoding HTT-interacting autophagy cargo receptor p62^47^ (**Figure S11B**).

Taken together, the GO analysis results indicate that HD-associated gene expression changes in human MSNs are distinct from those of other striatal neurons in HD, and point to the different cellular processes that are either induced or repressed at the level of transcription in MSNs. The large magnitude of HD-associated down-regulation seen for many genes (median log2FoldChange −0.67 and −0.73 for genes downregulated in dMSNs and iMSN, respectively) indicates that these changes likely occur in the majority of nuclei being assayed.

### DNA repair pathways

Given the expansions of *mHTT* CAG tract in MSNs, we investigated whether there are disease-associated changes in the transcript levels of genes encoding MMR and BER proteins. These data revealed clear distinctions in the regulation of these pathways in MSNs compared to interneurons. Notably, we found that in HD MSNs, *MSH2* and *MSH3* expression levels are significantly reduced relative to MSNs from control donors, while *POLD1*, coding for the large catalytic subunit of the DNA polymerase delta complex, undergoes a disease-associated upregulation that is not entirely specific to MSNs (**Figure 7C**). Notably, *POLD1* was recently identified as a candidate modifier of HD^49^. The expression of several DNA glycosylases (*NEIL1*, *UNG* and *TDG* genes) involved in the detection of DNA lesions is also changed in MSNs to a greater degree than in other neuronal cell types. Although further validation will be required to define and understand these changes, transcriptional regulation of DNA repair genes in response to genotoxic stress has been demonstrated in both rodent and human cells^50^. It seems likely, therefore, that further studies of these pathways will shed light on additional features of DNA repair that occur in concert with or as a result of *mHTT* CAG expansion.

### Genes required for MSN viability and functionality

A second important class of genes that we chose to analyze in more detail were those previously shown to preserve MSN viability in mice expressing *mHTT* with extra-long CAG tract^51^. Since decreased expression of these genes in HD MSNs could contribute to their loss, we asked whether any of these genes were strongly downregulated in HD MSNs. **Figure 7D and Figure S11C** depict genes that were found to be essential for MSN viability in the zQ175 and R6/2 mouse models, respectively, and had undergone the largest expression level decreases in human MSNs in HD donors. We consider these genes as candidates for further investigation as they can help to identify molecular processes that may contribute to loss of MSN viability in HD. Among these, HD-associated downregulation of TATA-binding protein-associated factor 1 gene (*TAF1*) in MSNs is an example of a change that could be detrimental to human MSN survival, as reduced expression and aberrant splicing of *TAF1* is thought to be the cause of MSN loss seen in X-linked dystonia-parkinsonism^52–54^.

We also noted that the transcript levels of MSN-enriched *PDE10A* and *ANO3* undergo large disease-associated decreases equivalent in magnitude to complete silencing of these genes in >45 % and >60% of the remaining MSNs, respectively (*ANO3* transcript log2FoldChange −1.35 and −0.86 for dMSNs and iMSNs, *PDE10A* transcript log2FoldChange −1.44 and −1.33 for dMSNs and iMSNs). As missense mutations in ANO3 are known to cause dystonia (https://omim.org/entry/615034) and PDE10A mutations are known to cause childhood-onset hyperkinetic movement disorders (https://omim.org/entry/616921, in some cases with striatal degeneration: https://omim.org/entry/616922), it is likely that these transcript level changes have a significant effect on the functionality of a large fraction of remaining MSNs in HD.

### Distinctions between mouse and human transcriptional responses in HD MSNs

While extensive transcriptional dysregulation observed in human MSNs in HD donor samples is generally consistent with the “transcriptionopathy” that has been reported in HD mouse models^55^, most of the HD-associated nuclear transcript levels changes seen by FANSseq are not recapitulated in HD mouse models for which MSN-specific RNAseq data is available (**Figures S13A-S13C**). Conversely, many individual genes and pathways reported as altered in mouse models of HD are not exhibiting an HD-associated change in human FANSseq data. In particular, human FANSseq data does not support a consistent decrease in MSN marker gene expression (**Figure S14A**) or the induction of genes involved in innate immune signaling (**Figure S14B**)^45^. Also, most of the genes with altered expression in the striatum of BAC-CAG mouse model are not undergoing a similar transcript level change in human MSNs (**Figure S14C**)^56^, and the derepression of PRC2-complex target genes that accompanies neurodegeneration in the striatum of MSN-specific Ezh1/Ezh2 double knockout mice is not observed for the majority of these genes expressed in human MSNs (**Figure S14D**)^57^. These discrepancies are perhaps not surprising since gene expression patterns across striatal cell types can differ between human and mouse, as exemplified by the expression of *HAP1* and several other genes encoding Huntingtin interacting proteins (**Figure S15**). Also, while we have not focused on the transcriptional events that are altered in cell types that do not undergo somatic expansion in the human striatum, it seems highly probable that these data will also differ significantly between HD and mouse models, given that these cell types carry pre-expanded *mHTT* alleles in these models. Although it is difficult to directly compare the human FANSseq data with transcriptional changes occurring in bulk RNAseq data from the striatum or MSN translational profiling (TRAP) data from HD mouse models, the apparent discrepancies between molecular events occurring in mouse model systems and human HD donor samples place a strong emphasis on the importance of human data when initiating focused studies of candidate mechanisms of HD pathogenesis.

## DISCUSSION

Here we have used FANS^18^ to isolate thousands of nuclei of each neural cell type of human caudate nucleus and putamen to generate deep, high resolution, cell type-specific transcriptional and *HTT* CAG repeat tract length-measurement data from control and HD donors. Our data are consistent with the two-component model of HD pathogenesis^17^, adding that somatic *mHTT* CAG expansions alone are not sufficient to explain cell-type vulnerability, and reveal several cell type-specific molecular features of the disease.

### Somatic expansion of *mHTT* CAG tract is not sufficient to cause cell loss in the human striatum

The most vulnerable cell types in the HD striatum are MSNs^4^. While both dMSNs and iMSNs are progressively lost during the progression of the disease, iMSNs that express dopamine receptor D2 and enkephalin are most vulnerable in early stages^29^. Striatal interneurons are relatively spared early in the disease^5,7^. In particular, although CHAT+ INs are clearly affected, as indicated by reduced choline acetyltransferase (CHAT) activity in histological sections, the persistent expression of acetylcholinesterase (ACHE) in these cells indicates that they do not die during the disease^6,58^.

To understand the relationships between somatic CAG expansion and cell loss, we have analyzed the length of *HTT* CAG tract in gDNA extracted from nuclei of striatal cell types. While the stability of the *mHTT* CAG tract in SST+, TAC3+ and, PVALB+ INs can explain their relative resilience in HD, our data also demonstrate that large expansions of the *mHTT* CAG tract are not sufficient for loss of CHAT+ INs in HD. Furthermore, data collected from dMSN and iMSN nuclei isolated from HD donors (N=13 total) establish that the extent of somatic CAG expansion is equivalent in these two MSN subtypes despite the increased vulnerability of iMSNs relative to dMSNs^29^. This is especially evident in data from carriers of reduced penetrance *mHTT* alleles where the loss of MSNs is minor. While our data support the hypothesis that somatic CAG expansion is a first step in disease progression that is necessary for the loss of neurons in HD, they indicate also that additional factors must be required. This conclusion is strongly supported by studies of the human cerebral cortex in HD (accompanying manuscript Pressl et al) demonstrating that extensive expansion of the *mHTT* CAG tract occurs in many deep layer pyramidal cell types despite very selective loss of a specific class of corticostriatal projection neurons. Finally, our observation that in cerebellum the *mHTT* CAG tract is unstable selectively in Purkinje cells is consistent with somatic expansion of the CAG tract being a prerequisite of mHTT toxicity since these neurons have been shown to be vulnerable in HD^32, 59, 60^.

### Striatal MSNs are prone to somatic CAG expansion

The preferential expansion of the *mATXN3* CAG tract we detect in MSN nuclei isolated from SCA3 donor samples indicates that these neurons have a general propensity to expand long CAG tracts, which is not limited to the *mHTT* locus. Our results are thus suggesting that there are *trans*-acting factors that are rate-limiting in the process of repeat expansion or stabilization, and that their level of expression or activity differs between MSNs and cell types where both *mHTT* and *mATXN3* CAG tracts are more stable. Our finding that levels of both *MSH2* and *MSH3* are higher in MSNs compared to other striatal cell types suggests that the actions of the MutSβ complex at the *mHTT* locus could be a rate-limiting step that contributes significantly to the selective vulnerability of MSNs. This idea is supported by previous reports showing that both *MSH2* and *MSH3* are required for *mHTT* CAG expansion in a mouse model of HD^15, 16, 37^ and naturally-occurring *Msh3* variants that alter the levels of MutSβ affect somatic CAG expansion in an HD mouse model^38^. The mechanism by which MutSβ is required for repeat expansion is unclear. We offer mechanistic insight to how elevated MutSβ could be promoting somatic CAG expansions by showing that an excess of MutSβ inhibits FAN1 nucleolytic excision of excess CAG slip-outs-thereby allowing these to be retained as somatic expansions. In contrast, MutSα, a complex of MSH2 with MSH6, does not affect FAN1 excision of slip-outs, consistent with a lack of involvement of MutSα in somatic expansions^15^. While both MutSα and MutSβ bind to heteroduplex DNAs, MutSβ has a very different binding mode, stronger affinity, and longer protein-loop DNA lifetimes than MutSα^61–63^, and hydroxy-radical footprinting of MutSβ on CAG slip-outs showing extensive protection of the slip-out^64, 65^. Excess levels of FAN1 could partly overcome the extended dwelling time of MutSβ on a CAG slip-out^61, 63^, resulting in slip-out excision so as to avoid retention of the excess repeats. Therefore, our results also present a mechanistic explanation to how genetic variants that increase FAN1 expression can delay onset of HD^39^. The way in which MutSβ, an expansion-driver and disease hastener, modulates the nuclease activity of FAN1, an expansion suppressor and disease delayer, suggest that poor and efficient slip-out excision can result in expanded and stable CAG repeats in brain cell types that have high and low MutSβ to FAN1 ratio, respectively. We do note that our observations on elevated levels of *MSH2* and *MSH3* in MSNs do not rule out the presence of other factors that favor the expansion of *mHTT* CAG repeat tract in MSNs and CHAT+ INs.

Two additional features of somatic expansion are also evident from our data. First, the degree of somatic expansion of the *mHTT* CAG tract in MSNs exceeds the expansion evident in PCs. Second, in MSNs, the expansion of the *mHTT* CAG tract is more extensive than the expansion of the *mATXN3* CAG tract. Notably, in HD donors carrying the fully penetrant *mHTT* alleles, about half of the remaining MSNs have *mHTT* CAG tract length increases of at least 21 CAG repeats, whereas very few tracts that gain that many repeats are evident at *mHTT* allele in Purkinje cells or at *mATXN3* allele in MSNs. These data demonstrate that both the locus at which the expansion occurs, as well as cell-intrinsic factors determine the rate of expansion. The characteristics of *mHTT* exon 1 that make its CAG tract especially prone to expansion in MSNs remain to be determined.

### HD-associated changes accompanying mHTT CAG expansions

The first conclusion we can draw is that the presence of a very long CAG repeat tract in RNA, or polyQ domain in any protein does not necessarily lead to cell loss in the human brain. In addition to the resilience of CHAT+ INs in the face of large expansions of the *mHTT* CAG tract (discussed above), this conclusion is supported by the observation that despite expansion of *mATXN3* CAG tract to over 80 CAG repeats, we did not see a clear decrease in the number of MSN nuclei isolated from SCA3 donor samples. The latter finding is not inconsistent with neuroanatomical studies that have reported MSN loss in SCA3 to be variable^66^. These results are reminiscent of the detailed studies of SCA1 mouse models demonstrating that the presence of Capicua sensitizes cerebellar Purkinje cells to mutant *Atxn1*^67^. While these considerations place additional emphasis on proteins that interact with mHTT *in vivo*^68^, delineation of the cell types and subcellular domains in which these interactions occur will be crucial for understanding their role in HD pathogenesis.

A second important conclusion that can be drawn from our data is that extensive transcriptional disturbances occur in the majority of MSNs prior to their demise in the HD striatum. While these data are not by themselves sufficient to identify mechanisms of toxicity that cause neuronal dysfunction and death in the second step of HD pathogenesis, they raise an important issue that remains to be resolved: do these transcriptional changes indicate that MSN functions are significantly compromised long before they are lost in the HD brain, or are these transcriptional events homeostatic compensatory responses that allow MSNs to operate relatively well for an extended time? The strong induction of TFEB, a central regulator of lysosomal function and autophagy, and of a variety of its downstream targets argues that human MSNs mount an important defense against the mHTT misfolding and extra-nuclear aggregation in HD. Transcriptional induction of genes involved in somatodendritic and synaptic function may also be indicative of homeostatic mechanisms that respond to problems in these important neuronal functions. However, these positive responses to mHTT must be weighed against the nuclear functions that are strongly decreased in HD MSNs, likely as a direct effect of the presence of mHTT in the nucleus, and the wide variety of additional transcriptional changes that occur in these very vulnerable neurons. The large expression changes in several genes that have been shown to be required for MSN viability in R6/2 and Q175 mouse models of HD^51^, and in genes with a clearly established link to human MSN functionality (*ANO3*) and survival (*TAF1, PDE10A*) point to the possibility that it is more than just a small fraction of remaining MSNs that have become dysfunctional in HD.

A third important point that arises from our observation that both the FANSseq and ATACseq data we have collected suggest that the *RRM2B* and *SYT9* genes, which can modify HD age at onset^13^, are expressed more highly in glia. Although these data will need to be verified by further experimentation, the elevated expression of these genes in microglia (*RRM2B*) and oligodendrocytes (*SYT9*) suggest that glia may contribute to HD pathogenesis. It is noteworthy that a major role for glial dysfunction has been proposed in a variety of other late-onset neurodegenerative disorders^69, 70^. However, even if the onset of motor symptoms in HD is modulated by the effect genetic variants in *RRM2B* and *SYT9* have on glial cells, our results suggest that the role of glia in HD progression is likely to be secondary given that the *mHTT* CAG tract instability is minimal in these cell types.

### Concluding remarks

The data we have presented here shed new light on several important features of HD pathogenesis. While we have highlighted some of the cell type-specific transcript level changes that are associated with the occurrence of somatic CAG expansion, future studies are needed to clarify the degree to which these changes perturb the implicated biological processes in human brain, and whether these changes are detrimental or compensatory. We hope that further analyses of the comprehensive datasets we have provided will stimulate others to interrogate them in the context of detailed mechanistic studies of HD. However, it is important to recognize that somatic expansion of *mHTT* CAG tract occurs only in select CNS cell types in HD, and that there are species-specific differences in cell type-specific gene expression in the striatum. Careful consideration of these findings will be important when studying HD molecular pathogenesis in models based on other species.

## ACKNOWLEDGEMENTS

This study was supported by funding from the CHDI Foundation, and we are thankful to Thomas F. Vogt and Jian Chen for helpful discussions throughout the project. We thank Nicholas Didkovsky, Cuidong Wang, Dr. Hasnahana Chetia and Yan Coulombe for technical assistance, Dr. Marc Ciosi for his advice on CAG tract sizing assay and Dr. Kärt Mätlik for advice on data visualization and for critical comments on the manuscript. We are also grateful to the Rockefeller University Genomics Resource Center for advice and support. We thank Jean Paul G. Vonsattel and Andrew F. Teich from Columbia University Alzheimer’s Disease Research Center (funded by NIH, grant P30AG066462), Dirk Keene from University of Washington BioRepository and Integrated Neuropathology Laboratory (supported by the Alzheimer’s Disease Research Center, grants AG066567 and AG066509), The University of Michigan Brain Bank (P30AG053760/ P30AG072931 University of Michigan Alzheimer’s Disease Core Center), Netherlands Brain Bank, Harvard Brain Tissue Resource Center and UCLA Human Brain & Spinal Fluid Resource Center (supported by National Institutes of Health and the US Department of Veterans Affairs) for assisting and providing postmortem brain tissues. K.M. was supported by a fellowship from the Sigrid Jusélius Foundation. A.L.D. is supported by a Postdoctoral Researcher Fellowship from the Hereditary Disease Foundation and the Fox Family Foundation. J.-Y.M. is supported by the Canadian Institutes of Health Research (FRN-388879). C.E.P. is supported by the Canadian Institutes of Health Research (FRN-148910; FRN-173282), the Natural Sciences and Engineering Research Council of Canada (RGPIN-2016-08355 RGPIN-2016-06355/498835), The Petroff Family Foundation, The Marigold Foundation, Tribute Communities, and the Fox Family Foundation. J.-Y.M. holds a Tier 1 Canada Research Chair in DNA Repair and Cancer Therapeutics. C.E.P. holds a Tier 1 Canada Research Chair in Disease-Associated Genome Instability.

## AUTHOR CONTRIBUTIONS

Conceptualization: K.M., C.E.P., N.H., Formal Analysis: K.M., M.R.P., T.C., A.L.D., Investigation: K.M., M.B., A.L.D., Resources: L.K., D.A.D., M.-C.C., J.-Y.M., N.H., Visualization: K.M., Writing - original draft: K.M., N.H., Writing - review and editing: M.R.P., M.B., L.K., C.E.P., Supervision: C.E.P., J.-Y.M., N.H., Funding acquisition: C.E.P., N.H.

## RESOURCE AVAILABILITY

### Lead Contact

Further information and requests for resources and reagents should be directed to the Lead Contact, Nathaniel Heintz.

### Materials Availability

This study did not generate new unique reagents.

### Data and Code Availability

All sequencing datasets generated as part of this study will be made publicly available in NCBI GEO under accession GSE227729 (https://www.ncbi.nlm.nih.gov/geo/query/acc.cgi?acc=GSE227729).

## EXPERIMENTAL MODEL AND SUBJECT DETAILS

### Human Samples

For this work, fresh frozen brain samples were obtained from Miami’s Brain Endowment Bank™, University of Washington BioRepository and Integrated Neuropathology Laboratory, Columbia University Alzheimer’s Disease Research Center, The University of Michigan Brain Bank and Netherlands Brain Bank, or through the NIH NeuroBioBank and sourced from either the Harvard Brain Tissue Resource Center (HBTRC) or the NIH Brain & Tissue Repository-California, Human Brain & Spinal Fluid Resource Center, VA West LA Medical Center, Los Angeles, California. Drug addiction and schizophrenia as well as clinical evidence of brain cancers were reasons for sample exclusion, while samples from donors with a history of other non-brain cancers and diabetes were accepted. Caudate nucleus, putamen and cerebellar vermis were used for isolation of nuclei. The brain regions used from each donor and their age, race, sex and post-mortem interval, are noted in **Table S1**. The table includes information about the Vonsattel grade, calculated CAP100 score ^78^, the number of uninterrupted CAG repeats in their HTT alleles as well as the sequence of the CAG tract and the adjacent CCG tract, as determined from CAG tract length measurement data. De-identified tissue samples analyzed in this study were determined to be exempt from Institutional Review Board (IRB) review according to 45 CFR 46.102 (f).

## METHOD DETAILS

### Isolation, labeling and sorting of glial cell nuclei

Nuclei were isolated as described previously ^18^. For the labeling of glial cell nuclei and cerebellar granule cells, the isolated nuclei were washed once with Homogenization buffer (0.25 M sucrose, 150 mM KCl, 5 mM MgCl2, 20 mM Tricine pH 7.8, 0.15 mM spermine, 0.5 mM spermidine, EDTA-free protease inhibitor cocktail, 1 mM DTT, 20 U/mL SUPERase-In RNase inhibitor (Thermofisher #AM2696), 40 U/mL RNasin ribonuclease inhibitor (Promega #N2515)). Each washing step constituted of resuspension of nuclei pellet followed by centrifugation (1000 g, 4 min, +4⁰C).

Resuspended nuclei were fixed in Homogenization buffer with 1% formaldehyde for 8 min at room temperature followed by quenching with 0.125 M glycine for 5 min. Following centrifugation, the nuclei were washed once with Wash buffer (PBS, 0.05% TritonX-100, 0.5% BSA, 20 U/mL Superase-In RNase Inhibitor, 40 U/mL RNasin ribonuclease inhibitor) and incubated at room temperature on a shaker in Wash buffer for permeabilization and blocking of unspecific binding. Nuclei were washed twice in Wash buffer without TritonX-100 and resuspended in 100 µl of 40% ethanol containing TrueBlack Lipofuscin Autofluorescence Quencher (Biotium, #23007) for 40-50 seconds. Nuclei were washed twice with Wash buffer (w/o TritonX-100) and incubated overnight at +4°C with the following antibodies: Rb x NeuN-Alexa647 (1:400, Abcam #ab190565), Rb x NeuN-Alexa594 (1:400, Abcam #ab207279), Mm x EAAT1 (1:2000, Santa Cruz Biotechnology, #sc-515839), Mm x IRF8-PE (1:65, ThermoFisher, #12-9852-82) and Goat x Olig2 (1:300, R&Dsystems, #AF2418). After three washes with Wash buffer (w/o TritonX-100) the nuclei were incubated for 30-45 min at room temperature with Donkey x Mm-Alexa488 (1:1000, ThermoFisher, # A-21202) and Donkey x Goat-Alexa647 (1:300, ThermoFisher, # A-21447). After three washes with Wash buffer (w/o TritonX-100) the nuclei were resuspended in Sorting buffer (PBS, 0.2% BSA, 40 U/mL RNasin ribonuclease inhibitor, 0.5 µg/mL DAPI). Aggregates of nuclei were excluded based on higher DAPI signal and the following gating strategies were used: neuronal nuclei (647+, 594+, 488-, large), oligodendrocyte nuclei (647+, 594-, 488-, small), microglia nuclei (647-, 594+, 488-, small) and astrocyte nuclei (647-, 594-, 488+, small). A separate sorting experiment was performed for collecting cerebellar granule cell nuclei. For this purpose nuclei were labeled with Rb x NeuN-Alexa594 (1:400, Abcam #ab207279) and Mm x ITPR1-Alexa488 (Santa Cruz Biotechnology # sc-271197 AF488), and granule cell nuclei were collected (488-, 594+).

For labeling neuronal nuclei, PrimeFlow labeling kit (Thermofisher, #88-18005-210) was used and fixation and permeabilization were carried out according to manufacturer’s instructions but with 200 U/mL Superase-In RNase inhibitor and 400 U/mL RNasin ribonuclease inhibitor present at every incubation step. For sorting the nuclei were resuspended in sorting buffer (PBS, 0.2% BSA, 40 U/mL RNasin ribonuclease inhibitor, 0.5 µg/mL DAPI). Probes specific to DRD1 (Alexa-647, # VA1-3002351-PF), DRD2 (Alexa-488, # VA4-3083767-PF) and PPP1R1B (Alexa-568, # VA10-3266354-PF) were used to label dMSN (647+, 568+, 488-, large) and iMSN nuclei (647-, 568+, 488+, large). In a separate set of experiments, probes specific to TAC3 (Alexa-647, # VA1-16603-PF), ETV1 (Alexa-488, # VA4-3083818-PF), SST (Alexa-568, # VA10-3252595-PF) and PPP1R1B (Alexa-568, # VA10-3266354-PF) were used to label the nuclei of TAC3+ interneurons (647+, 568-, 488+), PVALB+ interneurons (647-, 568-, 488+), SST+ interneurons (647-, 568+++, 488-) and MSNs (647-, 568+, 488-, large). Probes specific to TRPC3 (Alexa-647, # VA1-3004835-PF), COL6A6 (Alexa-647, # VA1-3014134-PF) and PPP1R1B (Alexa-568, # VA10-3266354-PF) were used in another set of experiments to label cholinergic interneuron nuclei (647+, 568, large) and MSN nuclei (647-, 568+, large). CA8 probe (Alexa-647, # VA1-3001892-PF) was used for sorting Purkinje neuron nuclei (647+, large). Aggregates of nuclei were always excluded based on higher intensity of DAPI staining. All PrimeFlow target probes were used at a dilution of 1:40.

### ATACseq library preparation

For generating ATACseq data the nuclei were treated with Tagment DNA TDE1 Enzyme (Illumina, #15027865) before fixation and labeling. The exact number of nuclei processed depended on the abundance of the population labeled and collected. Briefly, 800,000 nuclei were pelleted by centrifugation (5 min at 950xg) and resuspended in 10 mM Tris-HCl pH 7.6, 10 mM NaCl, 3 mM MgCl2, 0.01% NP-40 followed by centrifugation (500xg for 10 min at +4°C). The pellet was resuspended in 200uL of Transposition Mix (1x TD buffer containing 20 U/mL Superase-In RNase Inhibitor, 40 U/mL RNasin ribonuclease inhibitor and 1.25 µl of Illumina Tagment DNA TDE1 Enzyme per every 100 000 nuclei) and incubated at +37°C for 30 minutes. The reaction was stopped and nuclei fixed by adding 1mL of Homogenization buffer with 1mM EDTA and 1% formaldehyde. After 8 min of incubation on a shaker the fixative was quenched by adding glycine (0.125 M) for 5 min. After washing the nuclei once in Homogenization buffer and once in Wash buffer (w/o TritonX-100) the sample was processed like described above, proceeding with the steps that follow permeabilization of nuclei. After sorting, the collected nuclei were centrifuged at 1600xg for 10 min +4°C and resuspended in 200 uL of RC solution (50 mM Tris-HCl pH 7.6, 200 mM NaCl, 1 mM EDTA, 1% SDS, 5 µg/mL Proteinase K) and incubated overnight at 5512.Incubate at +55°C. Genomic DNA was isolated with using MinElute Reaction Cleanup Kit (Qiagen, #28206) and used for PCR amplification (72°C – 5min, 98°C – 30sec, 12-14x[98°C - 10sec, 63°C – 30sec, 72°C – 1 min]) with NEBNext® High-Fidelity 2X PCR Master Mix (New England Biolabs, #M0541S) and barcoded Nextera primers (1.25 μM each) ^22^. Following double-sided size selection by bead-purification the libraries were quantified with Qubit dsDNA HS assay kit (ThermoFisher #Q32851) and pooled for sequencing on NovaSeq6000 (SP 2 x 100bp).

### FANSseq library preparation and sequencing

RNA extraction was carried out with AllPrep DNA/RNA FFPE Kit (Qiagen, #80234) with modifications described previously (Xiao 2018). RNA-seq libraries were prepared with Trio RNA-Seq™ library preparation kit (Tecan, #0506-A01), quantified with Qubit dsDNA HS assay kit (ThermoFisher #Q32851) and pooled for sequencing on NovaSeq6000 (SP 2 x 150bp).

### RNAseq Data Processing

Sequence and transcript coordinates for human hg38 UCSC genome and gene models were retrieved from the BSgenome.Hsapiens.UCSC.hg38 Bioconductor package (version 1.4.1) and TxDb.Haspiens.UCSC.hg38.knownGene (version 3.4.0) Bioconductor libraries respectively.

FANSseq reads were aligned to the genome using Rsubread’s subjunc method (version 1.30.6) ^71^ and exported as bigWigs normalized to reads per million using the rtracklayer package (version 1.40.6). Reads in genes were counted using the featurecounts function within the Rsubread package against the full gene bodies (Genebody.Counts) and gene exons (Gene.Counts).

### ATACseq Data Processing

The ATACseq reads were aligned with the hg38 genome from the BSgenome.Hsapiens.UCSC.hg38 Bioconductor package (version 1.4.1) with Rsubread’s align method in paired-end mode. Fragments between 1 and 5,000 base pairs long were considered correctly paired. Normalized, fragment signal bigWigs were created with the rtracklayer package. Peak calls were made with MACS2 software in BAMPE mode ^72, 79^. For each striatal interneuron type except cholinergic interneurons, the ATACseq consensus peaks were called from 4 ATACseq datasets generated from four different control donors. For MSNs, ATACseq consensus peaks were called from 8 dMSNs datasets from 7 different HD donors, from 9 iMSN datasets from 8 different HD donors, and 31 dMSNs datasets and 32 iMSN datasets from 8 control donors (up to four datasets from each donor). High confidence consensus peaks were derived by creating a nonredundant peak set for each cell type and disease state, and then filtering down to peaks that were present in the majority of samples. These were then annotated to TSS based on proximity using the ChIPseeker package (version 1.28.3) ^73^. NCBI Refseq hg38 gene annotation was used (version 109.20211119).

### Differential gene expression analysis and principal components analysis

For comparison of transcript abundance data between different cell types from control donors, the comparisons of data from caudate nucleus and putamen were done independently. For control donors from which there was data available from both posterior and anterior parts of the same structure, a single table of average raw read counts per gene was generated for each cell type. For comparison of control donor data to HD donor data, up to four separate datasets for a given cell type (anterior putamen, posterior putamen, anterior caudate nucleus, posterior caudate nucleus) were combined by calculating the average raw read counts per gene (rounding up to integer), so that each donor was represented by a single FANSseq dataset for each cell type. Principal component analysis plots for 500 most variant genes were generated with pcaExplorer ^74^ using average raw read ‘Genebody.Counts’ tables as input data (one table for each donor per cell type). Average raw read ‘Gene.Counts’ tables (i.e. derived from FANSseq reads mapped to exons), one for each donor per cell type, were converted into normalized counts by DESeq2, thereby accounting for sequencing depth differences, and used for differential gene expression analysis by DESeq2^75^ ^80^ (version 1.36.0). Differential gene expression analysis performed based on ‘Genebody.Counts’ (i.e. FANSseq reads mapped to full gene bodies) is also provided. For the visualization of gene expression differences across cell types, ‘expression in cell type A’ was calculated as the mean of DESeq2-normalized counts from each donor. ‘Expression in a cell type A’ was then turned into relative expression (‘relative expression in cell type A’ = ‘expression in a cell type A’ divided by ‘mean of expression in all cell types compared’) and the resulting values were log2-transformed for visualization by Pheatmap R package (version 1.0.12). Relative expression was calculated in the same manner when comparing expression across individual samples instead of cell types. Differential gene expression analysis results were filtered to exclude genes for which none of their annotated TSS positions in NCBI Refseq hg38 (version 109.20211119) overlapped with ATACseq consensus peaks defined separately for the cell types compared. These lists were augmented with a small number of genes (<110) for which manual inspection of mapped FANSseq and ATACseq reads in Integrative Genomics Viewer ^76^ suggested that these genes were in fact expressed.

### Pathway enrichment analysis

The filtered lists of differentially expressed genes (DEGs) (p adj < 0.05) with accessible TSS regions were used for analysis of over-representation of Gene Ontology Cellular Compartment (GOCC) terms with clusterProfiler package ^77^ (version 4.4.4, GOSOURCEDATE: 2022-03-10). The augmented list of all genes with accessible TSS regions was used as the ‘background’ list for comparison (‘universe’), and the following parameters were used: qvalueCutoff = 0.05, minGSSize = 5, maxGSSize = 2000. For identifying GOCC pathways over-represented among genes that showed disease-associated up- or downregulation only in dMSNs, the list of DEGs from ‘HD_dMSN vs ctrl_dMSN’ comparison (P adj. < 0.05) was filtered to exclude genes that had changed expression (P adj. < 0.05) in the same direction in any of the three ‘HD_interneuron vs ctrl_interneuron’ comparisons (TAC3+ INs, PVALB+ INs or SST+ INs).

### *HTT* and *ATXN3* CAG tract sizing

Genomic DNA was purified using AllPrep DNA/RNA FFPE Kit (Qiagen, #80234) and concentrated in a vacuum concentrator if required. *HTT* CAG tract sizing was done by next generation sequencing of PCR amplicons of *HTT* exon1 using a modified version of a previously published protocol ^28^. Up to 10 ng of gDNA was amplified in a 20 μl volume using NEBNext® High-Fidelity 2X PCR Master Mix (New England Biolabs, #M0541S) supplemented with 5% dimethyl sulfoxide and barcoded primers specific to *HTT* Exon 1 (0.5 μM each) ^28^ or *ATXN3* exon 10: 1 cycle 96°C - 5 min, 30x [96°C - 45s, 61°C - 45s, 72°C - 3min], 72°C - 10min. The number of amplification cycles was raised to 32 cycles if the amount of gDNA input was below 4 ng. After PCR the samples were combined into small pool of 2-6 samples and size selection was carried out by adding 0.55x volume of AMPure XP beads (Beckman Coulter, #A68831). The concentrations of purified library pools were quantified with Collibri™ Library Quantification Kit (ThermoFisher, #A38524100), combined into a sequencing library and sequenced on MiSeq sequencer using a 500 cycle MiSeq Reagent Nano Kit v2 with both index reads, but with 400 nt long read 1 and no read 2. Demultiplexed sequencing read data was aligned using Burrows-Wheeler Aligner (https://github.com/lh3/bwa, using BWA MEM default settings except: -O 6,6 -E 4,4) to a set of *HTT* exon 1 or *ATXN3* exon 10 reference sequences [suppl files 10 and 11] that differed by the number of CAG repeat units in the repeat tract. The number of reads uniquely mapped to each of the reference sequences in the set was considered to reflect the distribution of CAG tract lengths in the two *HTT* or *ATXN3* alleles in the cell population analyzed. *HTT* read mapping data from each donor was inspected manually for determining the nucleotide sequence of the adjacent polyproline tract and the presence/absence of interruptions in CAG tract. If *mHTT* exon 1 structure of the donor was atypical, then sequencing reads were realigned to a set of reference sequences matching that *mHTT* exon 1 structure. The length of CAG repeat tracts reliably mapped was limited to 113 repeat units. Uninterrupted CAG tract lengths of progenitor/unexpanded *mHTT* allele (M repeat units) and normal *HTT* allele (N repeat units) were defined from the two modes of mapped read length-distribution in CAG-sizing data from non-expanding cell types (usually striatal microglia and astrocytes, or, if available, cerebellar granule cells). R is the number of reads mapped to a reference sequence with the specified CAG tract length. The ratio of somatic expansions (RoSE) ^28^ and frequency of *mHTT* copies expanded by more than 20 repeat units (RU) were calculated as follows:

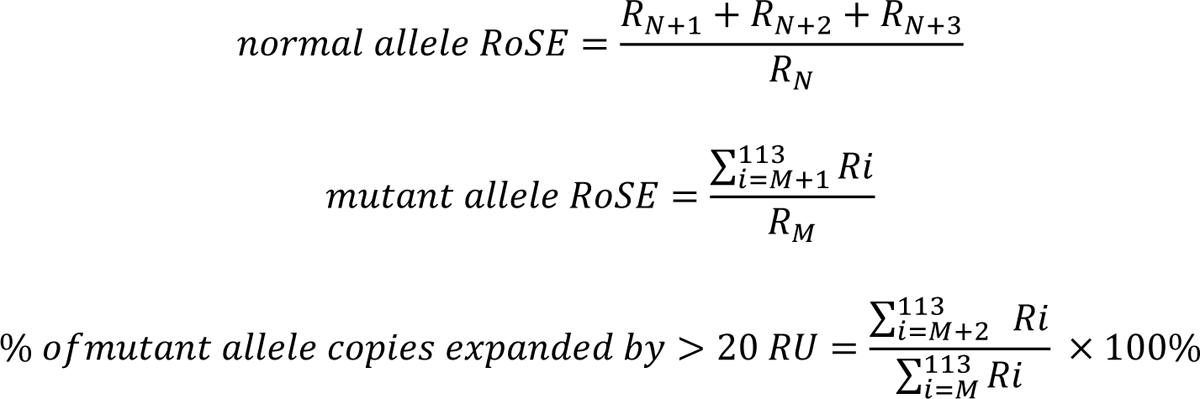

Quantification of CAG tract length changes for *mATXN3* was done in the same way. Statistical analysis of differences between cell types was carried out by comparing their ratio of somatic expansions or frequency of *mHTT* copies expanded by > 20 RU with one-way ANOVA, followed by Holm-Sidak’s multiple comparisons test.

### Western blotting

When isolating nuclei for western blotting the tissue homogenization and ultracentrifugation steps were carried out as described by Xiao et al. ^18^. After washing the nuclei once in Homogenization buffer the nuclei were resuspended in 1 mL of 1x PBS, 0.05% Triton X-100, 2% BSA and incubated at room temperature on a shaker for ∼ 15-20 minutes. The nuclei were labeled by adding the following antibodies: Rb x NeuN-Alexa647 (1:300, Abcam #ab190565), Rb x NeuN-Alexa594 (1:300, Abcam #ab207279), Mm x EAAT1-Alexa488 (1:200, Santa Cruz Biotechnology, #sc-515839 AF488), Mm x IRF8-PE (1:65, ThermoFisher, #12-9852-82) and Goat x Olig2 (1:200, R&Dsystems, #AF2418). After two washes with WB Wash buffer (1x PBS, 0.05% Triton X-100, 0.2% BSA) the nuclei were incubated for 30 min at room temperature with Donkey x Goat-Alexa647 (1:400, ThermoFisher, # A-21447). Nuclei were washed twice with WB Wash buffer and resuspended in Sorting buffer (w/o RNase inhibitors). After sorting, the collected nuclei were centrifuged at 1600xg for 10 min +4°C and the residual volume was kept to a minimum. Nuclei were treated with DNase I (in the presence of 0.5 mM Mg Cl2) at +37°C for 10 min, mixed with NuPAGE Sample Reducing Agent (ThermoFisher #NP0004) and β-mercaptoethanol (final conc. 4%, Sigma, #M3148), and heat-denatured at +96°C for 3 minutes. Material from 25 000 to 50 000 nuclei were loaded on NuPAGE™ 4 to 12% Bis-Tris Mini gels (ThermoFisher, #NP0322BOX), aiming for equal loading in each well. After blotting the samples onto nitrocellulose membrane and blocking unspecific binding by incubating the membrane in a 5% solution of non-fat dry milk, the membranes were probed with Rb x Histone H3 antibody (1:5000, Abcam, #ab1791) and Mm x Human MSH2 (1:300 BD Biosciences, #556349) or Mm x Human MSH3 (1:300 BD Biosciences, #611390) by incubating overnight at +4°C. After three washes with TBS-T (1x TBS, 0.1% Tween-20) the membranes were probed with IRDye® 680LT Donkey anti-Rabbit IgG Secondary Antibody (1:10 000, LICOR, #926-68023) and IRDye® 800CW Goat anti-Mouse IgG Secondary Antibody (1:10 000, LICOR, #926-32210) by 1 hour at room temperature. After three washes with TBS-T the membranes were imaged with Odyssey® DLx Imaging System.

### Protein purification

Recombinant human FAN1 protein was expressed from a Baculovirus and purified from Sf9 insect cells as described previously ^44, 81^. Recombinant human MutSα (MSH2-MSH6) and MutSβ (MSH2-MSH3) were generated from Sf9 insect cells using Baculoviruses expressing his-tagged hMSH2, hMSH3 and hMSH6, and a purification procedure described previously ^82–84^.

### FAN1 nuclease assay

FAN1 nuclease assays were performed as described ^44^ in nuclease assay buffer (50 mM Tris HCl pH 8.0, 25 mM NaCl, 1 mM MnCl_2_, 1 mM dithiothreitol, 200 mg/ml BSA) with 100 nM of fluorescently labeled DNA incubated with 50 nM of recombinant human FAN1 protein. Reactions were initiated by the addition of FAN1 protein, incubated at 37°C, for 20 minutes then stopped with formamide loading buffer (95% formamide, 10 mM EDTA). Reaction products were electrophoretically resolved on 6% denaturing sequencing gel for 1 hour at 2000 V and detected at fluorescence filter in the Typhoon FLA (GE Healthcare). Nuclease activity quantification compared the densitometric intensity of cleaved versus un-cleaved DNA (ImageQuant) In some of the experiment’s different incubation time and concentration of proteins are used and are mentioned in respective figure legends.

## SUPPLEMENTAL INFORMATION

List supplemental files:

Table S1 - List of samples used from each donor

## SUPPLEMENTAL FIGURES

**Figure S1.**
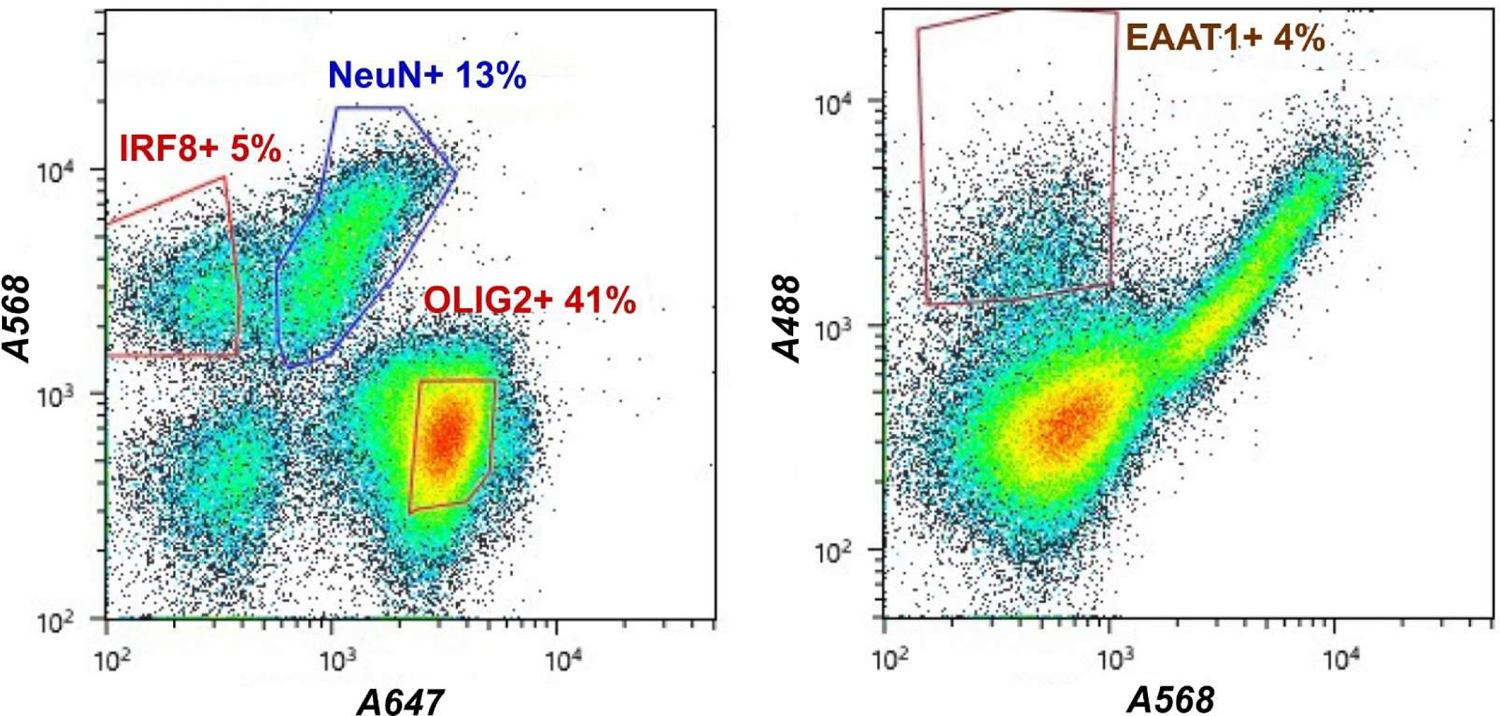
Representative FANS plots showing antibody-based labeling of glial cell nuclei isolated from human post-mortem caudate nucleus and putamen. The detailed strategy used for sorting is described in Materials and Methods.

**Figure S2.**
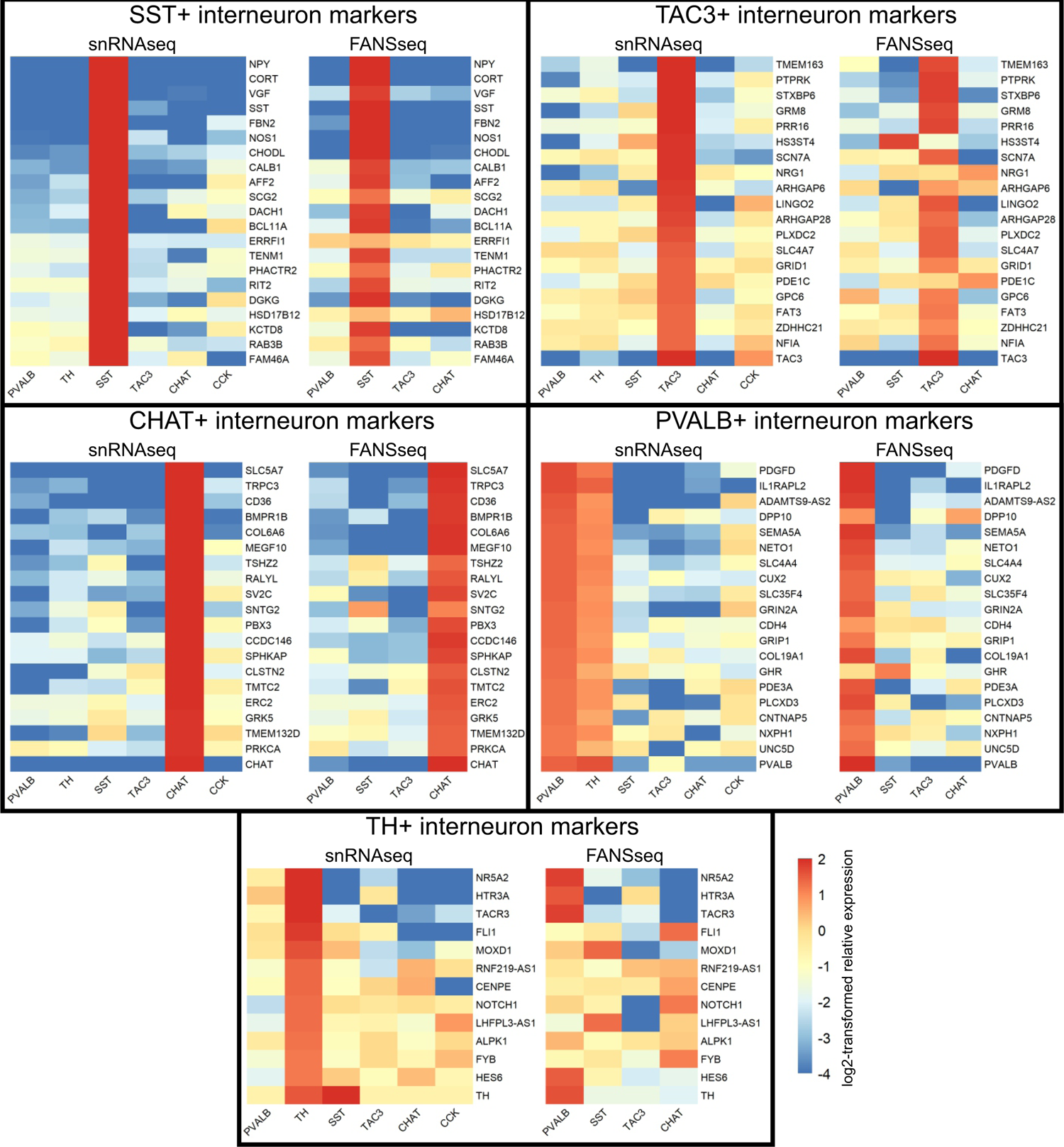
Comparison of interneuron populations collected using FANS to cell types defined from single-nucleus RNA sequencing of human striatum^1^. Relative expression in each cell type was calculated based on DESeq2-normalized counts from 6-8 control donors, and was log2-transformed for visualization (see Materials and Methods). The marker genes specific for each interneuron subtype were selected based on single-nucleus RNA sequencing data. The FANS-isolated ETV1+ TAC3-population of Parvalbumin-expressing interneuron nuclei most likely captured both the major PVALB+ interneuron population and the related smaller PVALB+ TH+ interneuron population defined as a separate subtype in snRNA-seq analysis^1^.

**Figure S3.**
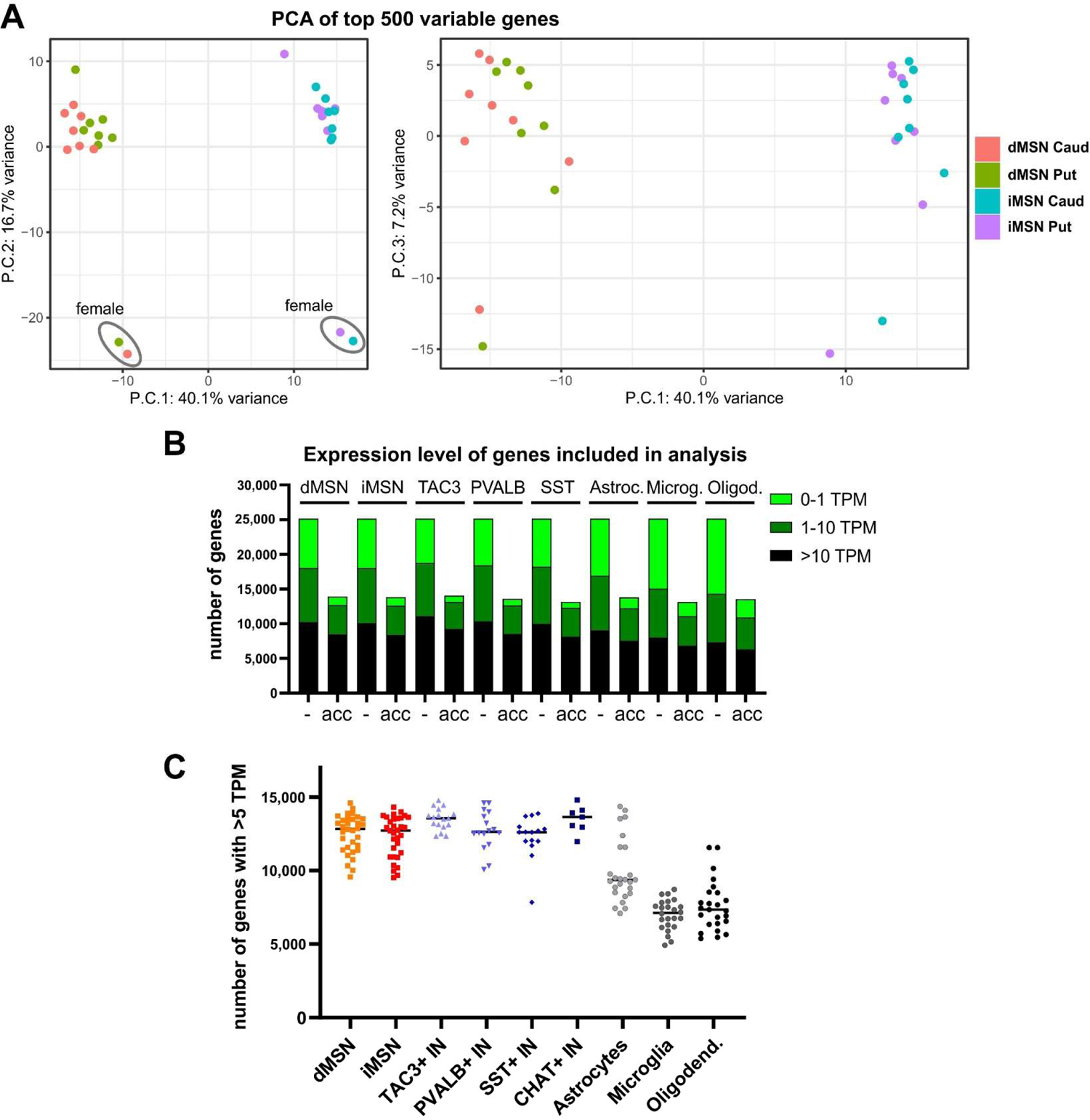
(**A**) Principal component analysis of FANSseq data from MSNs of caudate nucleus and putamen from 8 control donors. (**B**) The number of genes with accessible transcriptional start sites (acc.), defined as having a consensus ATACseq peak at the TSS, in striatal cell types from control donors. Genes have been grouped according to average expression level in FANSseq datasets from the specified cell type (TPM - transcripts per million). Excluding genes with inaccessible promoters removed a large proportion of genes with very low expression levels (< 1 TPM) while affecting fewer genes with moderate to high expression (>10 TPM). (**C**) Number of genes with expression level above 5 TPM in individual FANSseq datasets from control donors.

**Figure S4.**
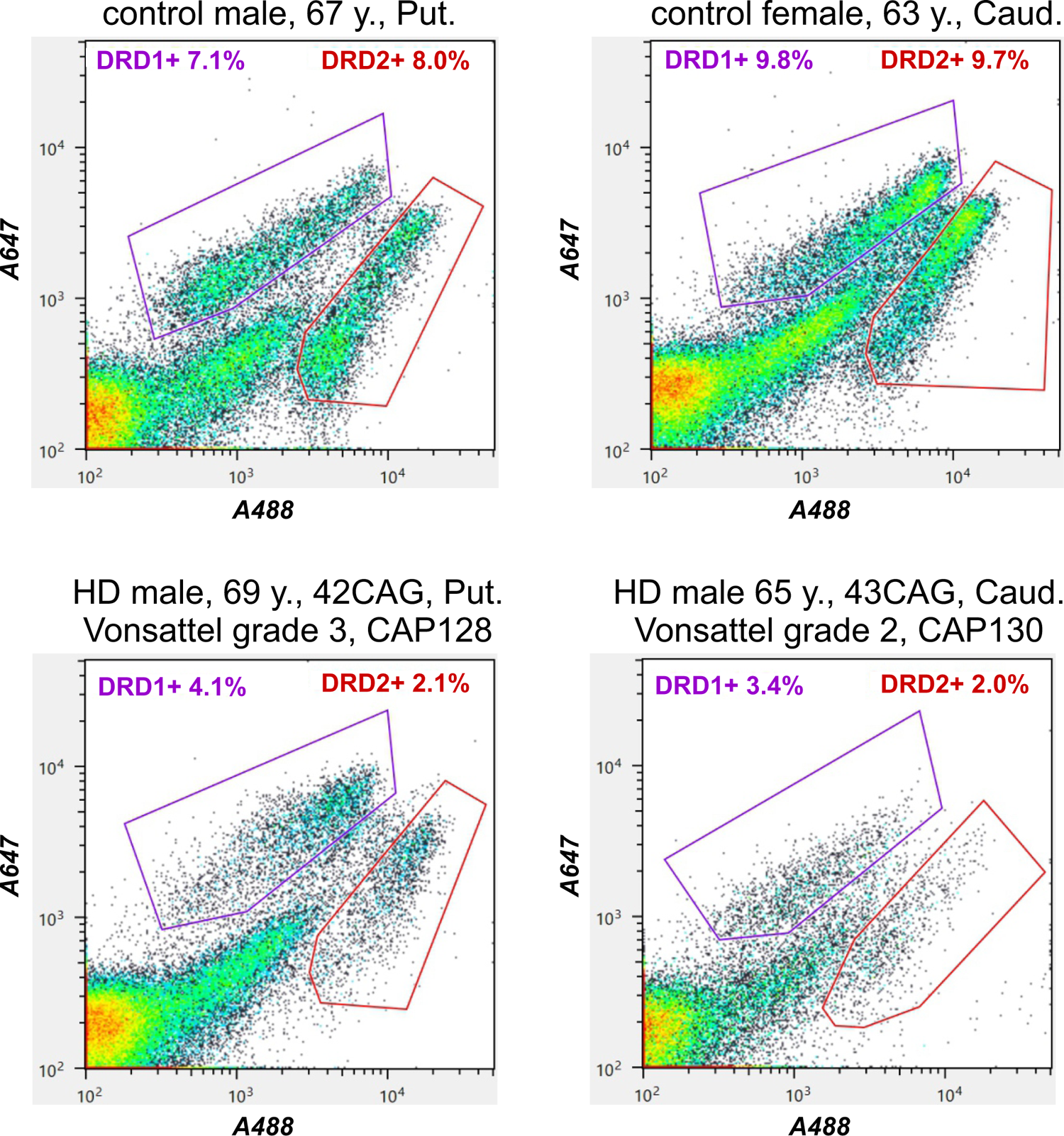
Representative FANS plots showing the labeling of dMSN and iMSN nuclei with Primeflow probes specific for *DRD1* and *DRD2* transcripts, respectively. Note that while there is a clear reduction in the abundance of MSN nuclei in striatal tissue from HD donors, these probes can still be used for the separation and isolation of MSN subtypes.

**Figure S5.**
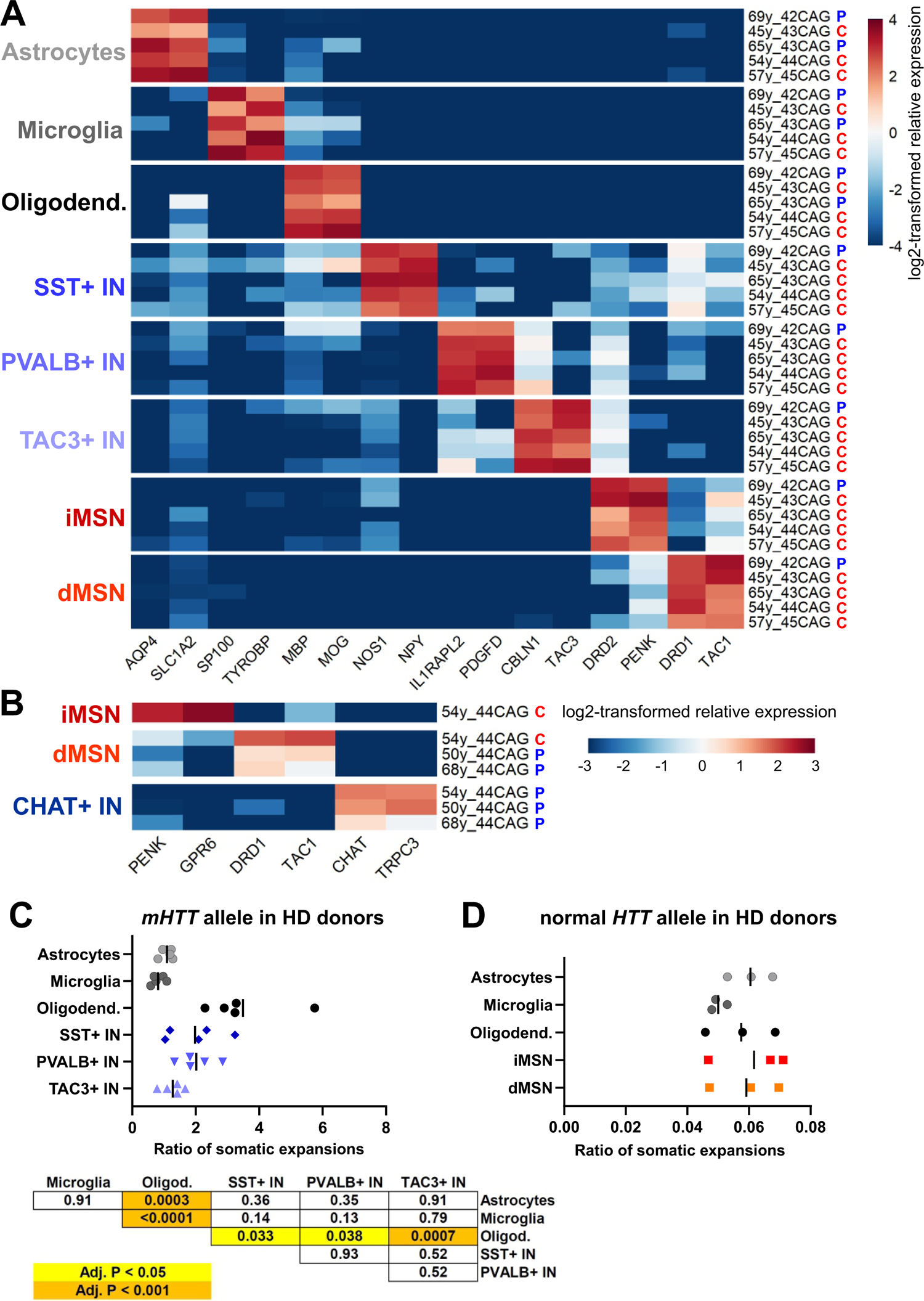
(A and B) Relative expression level of marker genes of striatal cell types in the nuclear transcriptome of nuclei collected for *mHTT* CAG repeat tract analysis. Heatmap depicts log2-transformed relative expression in each sample (calculated based on DESeq2-normalized counts). The striatal region of origin is indicated by colored letters (blue P – putamen, red C – Caudate nucleus). (C) Comparison of the calculated ratio of somatic expansions of *mHTT* CAG tract in striatal cell types other than MSNs. The table presents adjusted P-values as calculated by Holm-Sidak’s multiple comparisons test post one-way ANOVA (P < 0.0001). (D) Ratio of somatic expansions of normal *HTT* allele CAG tract in striatal cell types isolated from HD donors with 21-25 uninterrupted repeats in normal HTT allele. P = 0.599 according to one-way ANOVA.

**Figure S6:**
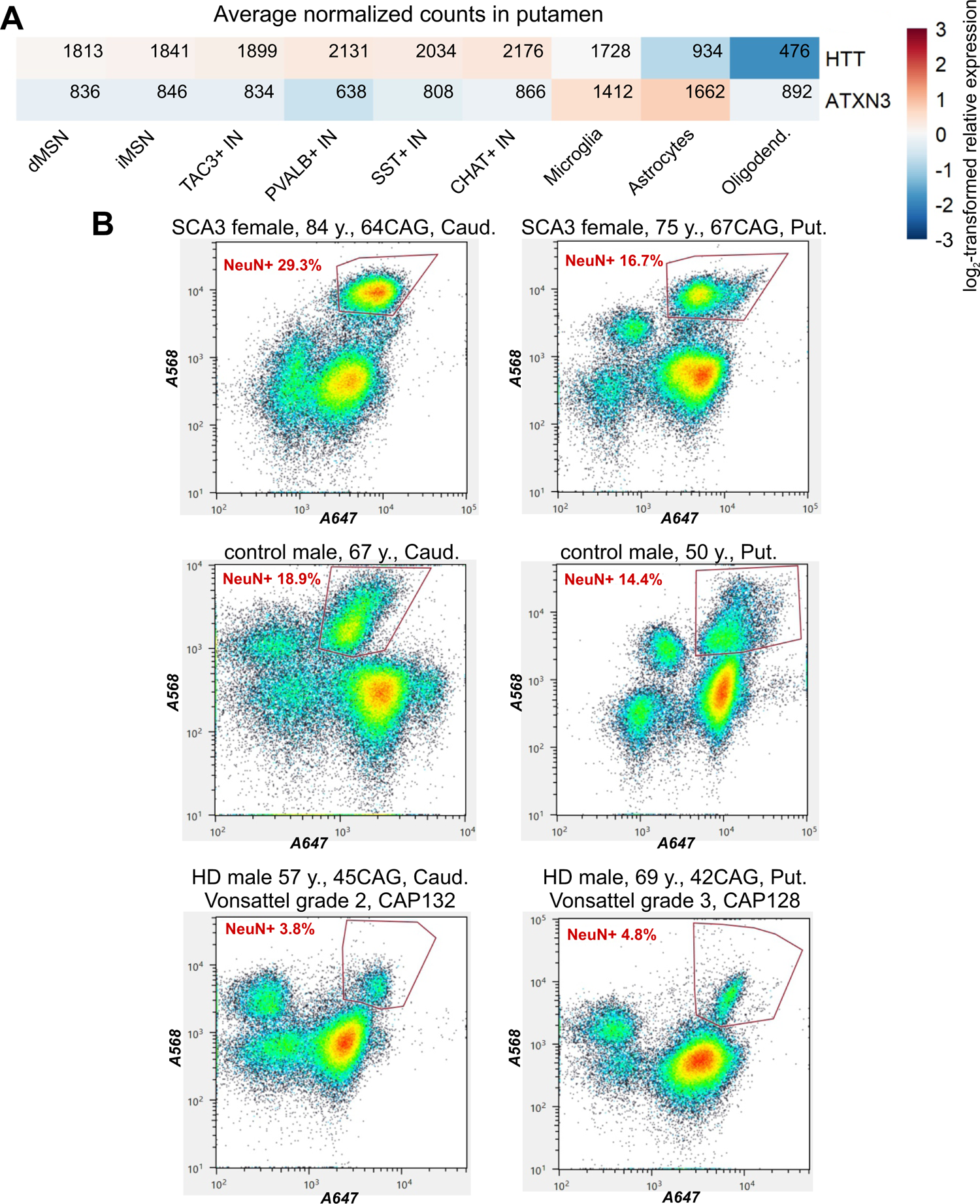
(**A**) Relative expression level of *HTT* and *ATXN3* in cell types of the putamen. Heatmaps depict log2-transformed relative expression in each cell type, calculated based on the mean of DESeq2-normalized counts from 6-8 control donors. (**B**) FANS plots showing the percentage of nuclei stained with anti-NeuN antibody in two oldest SCA3 donors, and in two representative HD donors and control donors.

**Figure S7:**
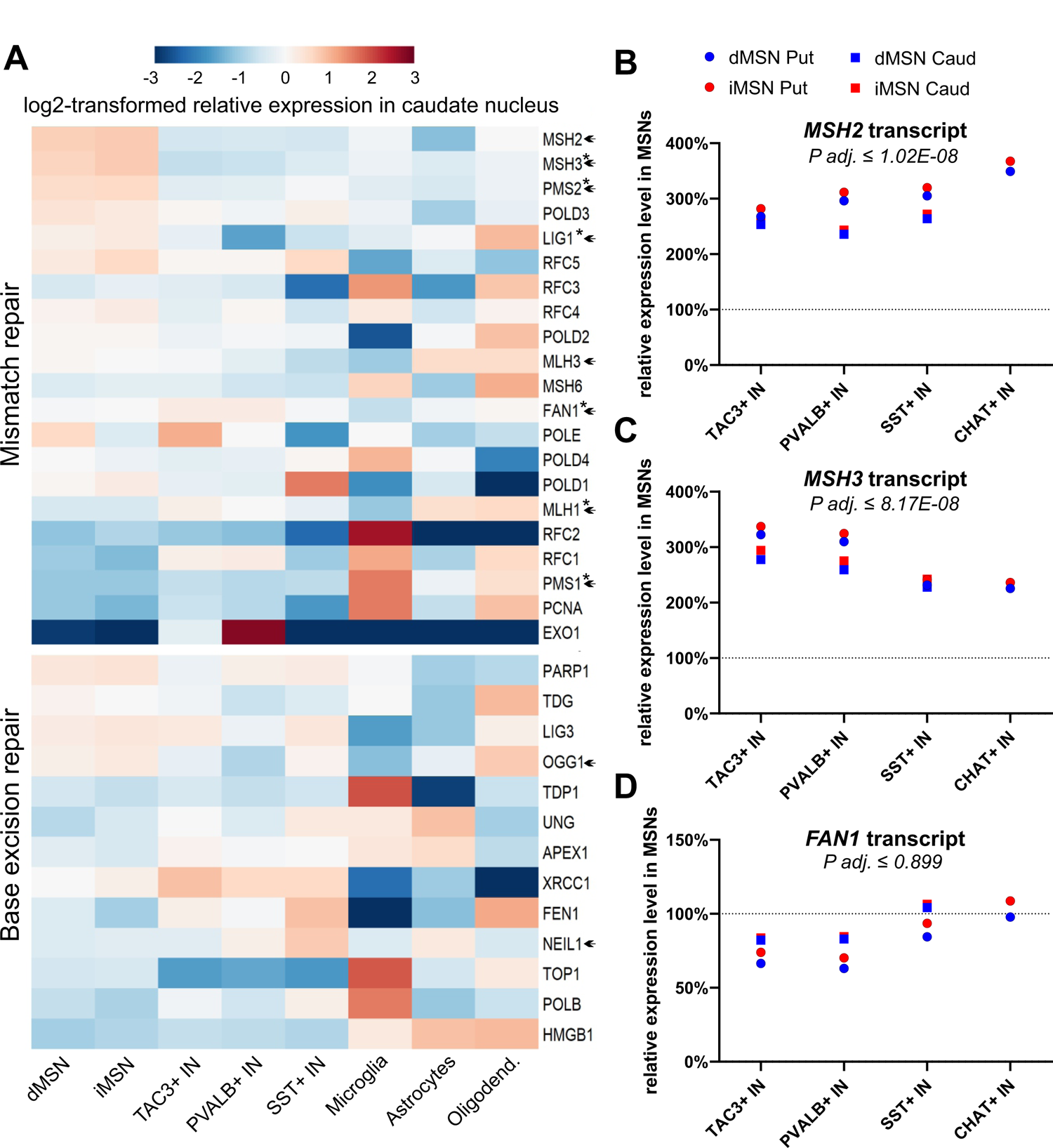
(**A**) Relative expression level of MMR and BER genes in cell types of caudate nucleus. Heatmaps depict log2-transformed relative expression in each cell type, calculated based on the mean of DESeq2-normalized counts from 7-8 control donors. Genes identified as HD age at onset-modifying candidates are marked with an asterisk. Genes known to influence CAG tract instability in HD mouse models are marked with an arrowhead. (**B-D**) Relative levels of (**B**) *MSH2*, (**C**) *MSH3* and (**D**) *FAN1* transcripts in MSNs compared to interneurons, calculated with DESeq2 using cell type-specific FANSseq data from control donors. The largest P adj. value of all MSN vs interneuron comparisons (by DESeq2) is indicated for each gene.

**Figure S8.**
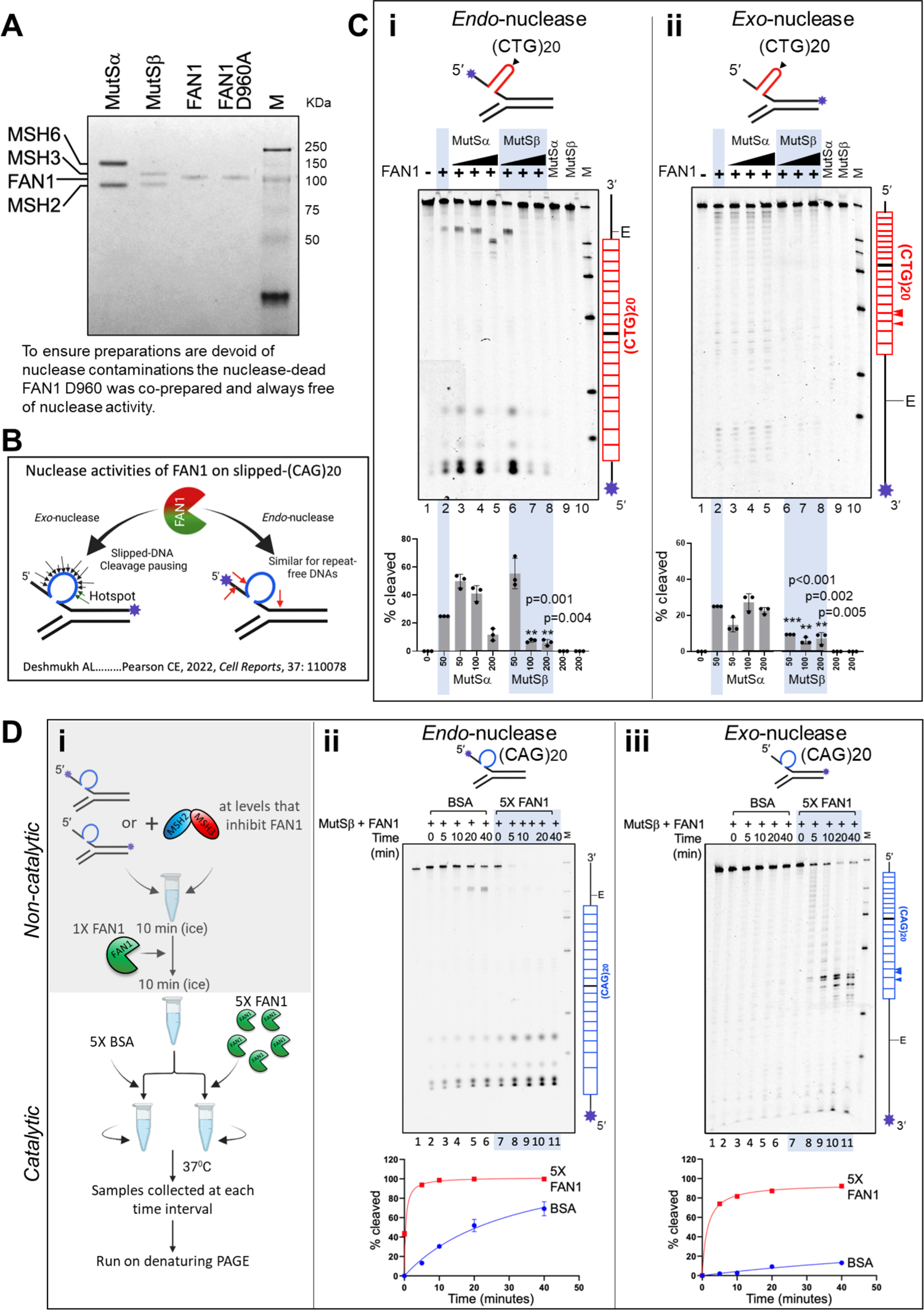
MSH2, MSH3, MSH6, and FAN1 proteins, nuclease assay, (CTG)20 digestion. (**A**) Coomassie brilliant blue-stained SDS-PAGE of purified human MutSα (MSH2-MSH6), MutSβ (MSH2-MSH3), FAN1 and nuclease-dead FAN1^p.D960A^ (purified in parallel to ensure an absence of contaminating nuclease activity in preparations), expressed in Sf9 cells using baculoviral overexpression. Both MutSα and MutSβ preparations were free of nuclease contamination (see panel Ci and Cii, lanes 9 and 10). Purity and activity were consistent for multiple preparations. (**B**) Schematic of FAN1 *exo*-nucleolytic and *endo*-nucleolytic cleavage sites on slipped-(CAG)20 DNA substrates labeled at the 3’ or 5’ ends, respectively. FAN1’s *exo*-nucleolytic ‘‘nibbling’’ of excess repeats parallels the ‘‘inchworm’’ expansions in patient brains, suggesting a role for FAN1 in regulating repeat instability^2^. *Endo*- and *exo*-nucleolytic activities can be distinguished by fluorescein amidite (FAM)-labeling at 5’ or 3’ ends of the (CAG)20 strand, respectively (asterisk); in this manner, only the labeled strand and it’s digestion products are tracked. (**C**) Slipped-(CTG)20 DNAs are digested by *endo*- and *exo*-nucleolytic activities of FAN1, which can be inhibited by MutSβ, but not by MutSα. FAM-labelled CTG-slip-out oligonucleotides (schematic, arrowhead indicated center of repeat tract) mimic intermediates of expansion mutations. *Exo*- and *endo*-nuclease activities were determined by labeling either 3′- or 5′-end of (CTG)20 strand, respectively (asterisk). “E” is elbow at the dsDNA-ssDNA junction. Slip-out DNA substrates (100 nM) were pre-incubated with increasing amounts of purified human MutSα or MutSβ (0-200 nM), and nuclease digestion was initiated by addition purified human FAN1 (50 nM). Vertical schematic to the right of each gel indicates location of cleavage sites along the FAM-labeled DNA strand. Percentage cleavage (densitometry) for each reaction was then normalized to cleavage levels for FAN1 alone, and these levels were graphed. N = 3 replicates. (**D**) Excess FAN1 overcomes MutSβ-mediated inhibition. (**i**) Flowchart outlining the competition experiment. Human MutSβ (200 nM), FAN1 (50 nM) and (**ii**) 5’- or (**iii**) 3′-FAM-labeled CAG-slip-outs (100 nM), i.e. conditions established to be completely inhibited by MutSβ, were preincubated in non-catalytic conditions (on ice). Reactions were split at the 10-min time point and challenged with 5X BSA (lanes 2–6) or an excess (5X) of FAN1 (lanes 7–11). Aliquots were taken for analysis at different timepoints (0, 5, 10, 20, & 40 minutes) and cleavage products were quantified (N=3).

**Figure S9.**
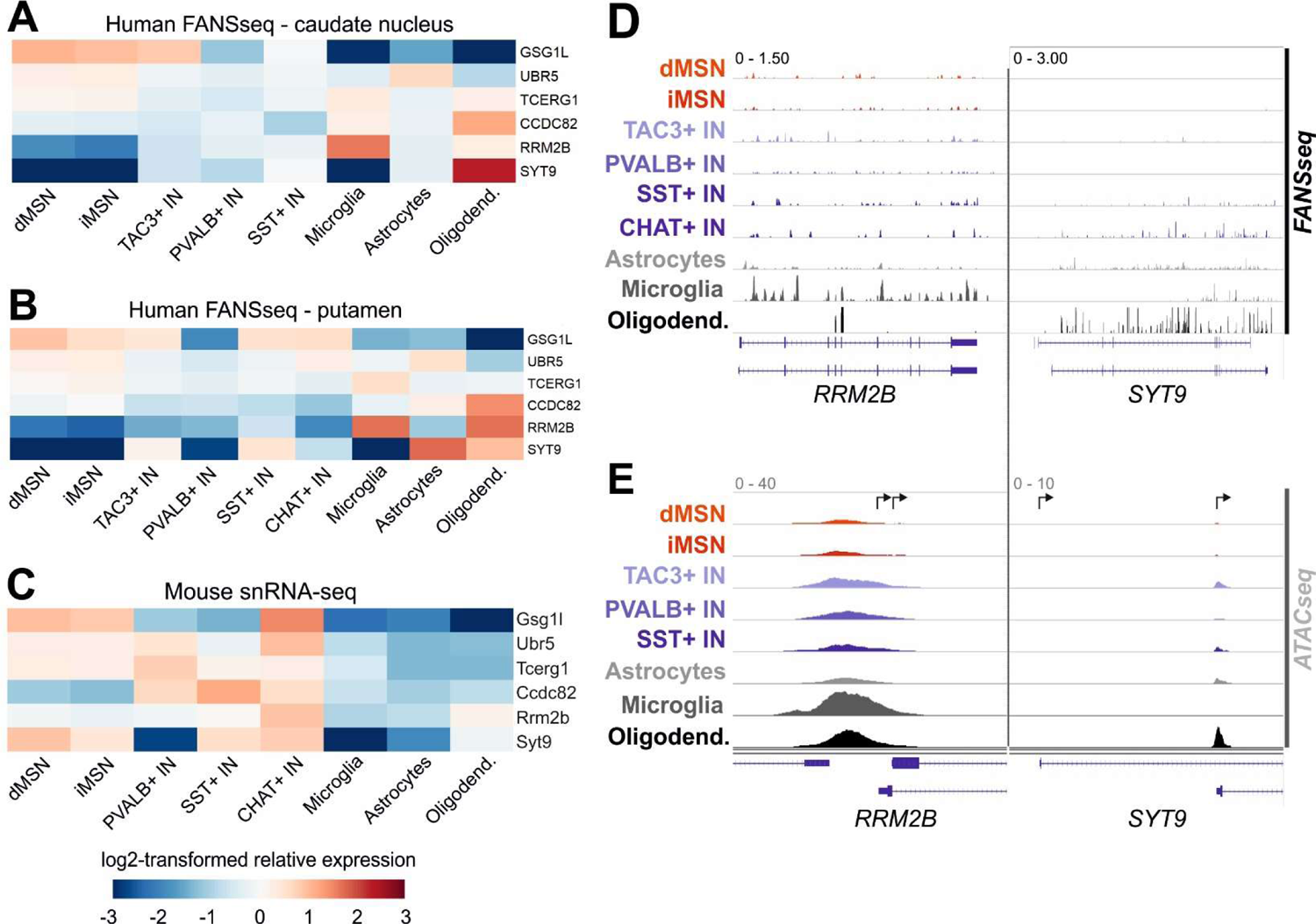
(**A and B**) Relative expression level of HD age-at-onset modifying candidate genes unrelated to DNA repair. Heatmaps depict log2-transformed relative expression in each cell type of (**A**) caudate nucleus and (**B**) putamen, calculated based on the mean of DESeq2-normalized counts from 6-8 control donors. (**C**) Cell type-specific expression according to snRNA-seq data from mouse striatum^3^, calculated from ‘transcripts per 100,000’ values. (**D**) Representative distribution of human FANSseq and (**E**) ATACseq reads mapped to *RRM2B* and *SYT9* genes. Arrows mark the position of annotated transcriptional start sites. The data are from a 41-year-old male control donor.

**Figure S10.**
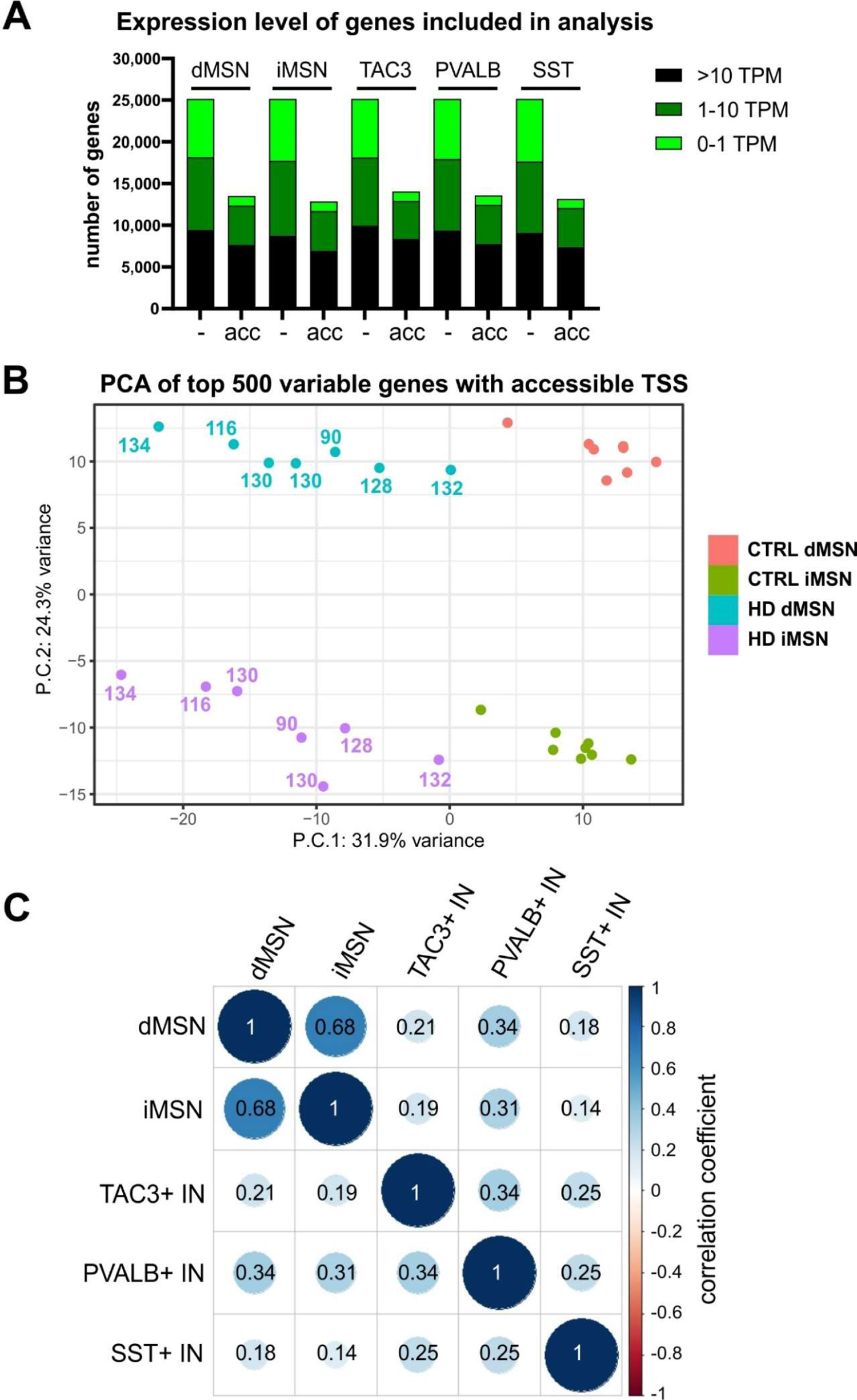
(**A**) The number of genes with accessible transcriptional start sites (acc.), defined based on the overlap of TSS and consensus peak of ATACseq data from HD donor MSNs and control donor interneurons. Genes have been grouped according to their average expression level in FANSseq datasets from HD donors (TPM - transcripts per million). **(B**) Principal component analysis of FANSseq data from putamen and caudate nucleus MSNs from HD and control donors, performed after the exclusion of genes with inaccessible TSSs. The calculated CAP100 scores are shown for each HD donor. **(C)** Correlation analysis of disease-associated expression changes of genes expressed in striatal neuronal types studied.

**Figure S11.**
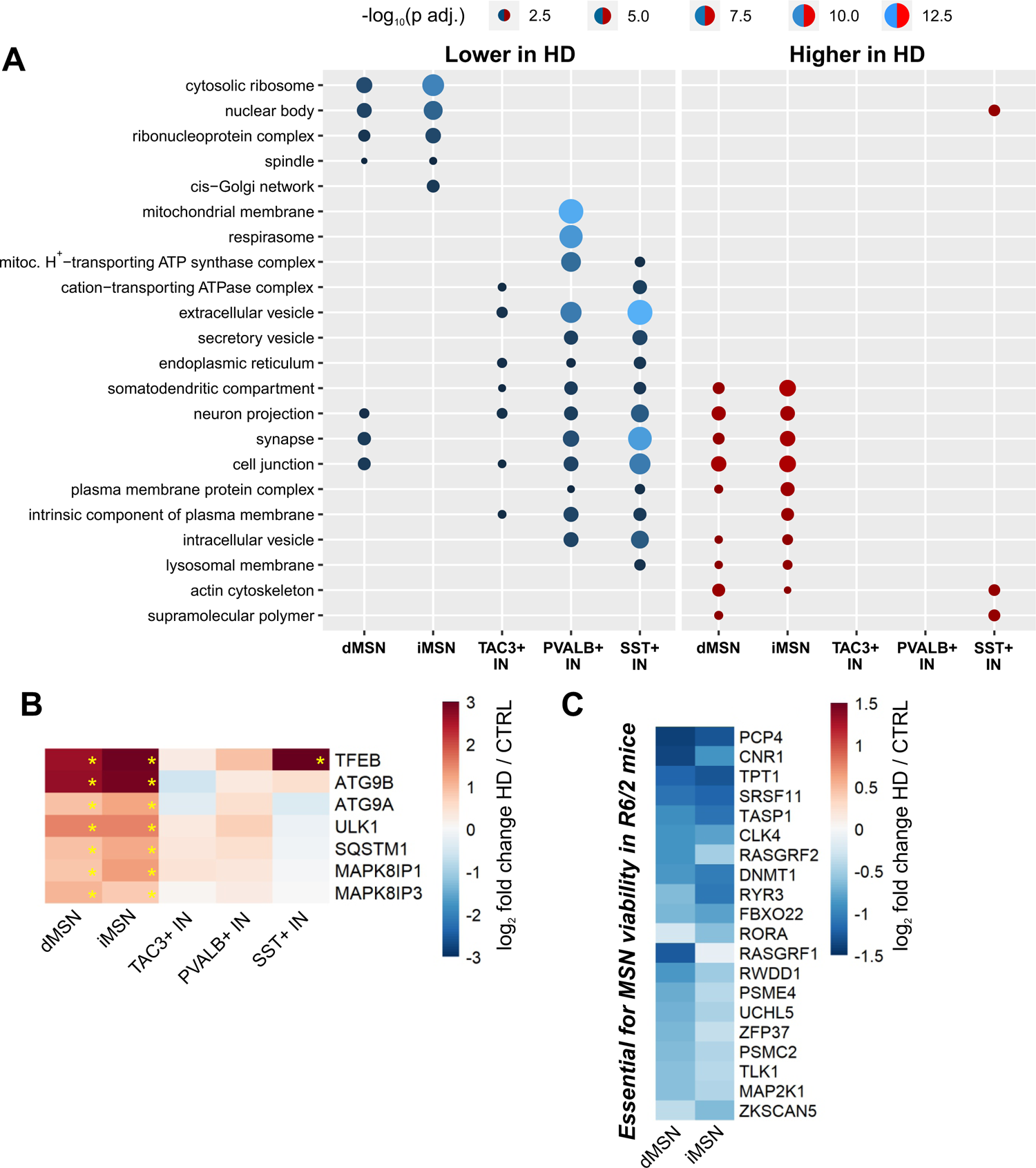
(A) Selected non-redundant Gene Ontology Cellular Component terms from enrichment analysis of genes with disease-associated expression changes in each striatal neuron type (p adj < 0.05). (B) Heatmap depicting disease-associated changes in transcript levels of selected genes regulating autophagosome formation and transport. Asterisks denote p adj < 0.05. (C) Genes essential for MSN viability in the R6/2 mouse model of HD^4^ that are also downregulated in HD iMSNs or dMSNs by more than a third (log2FC<-0.6, p<0.01).

**Figure S12.**
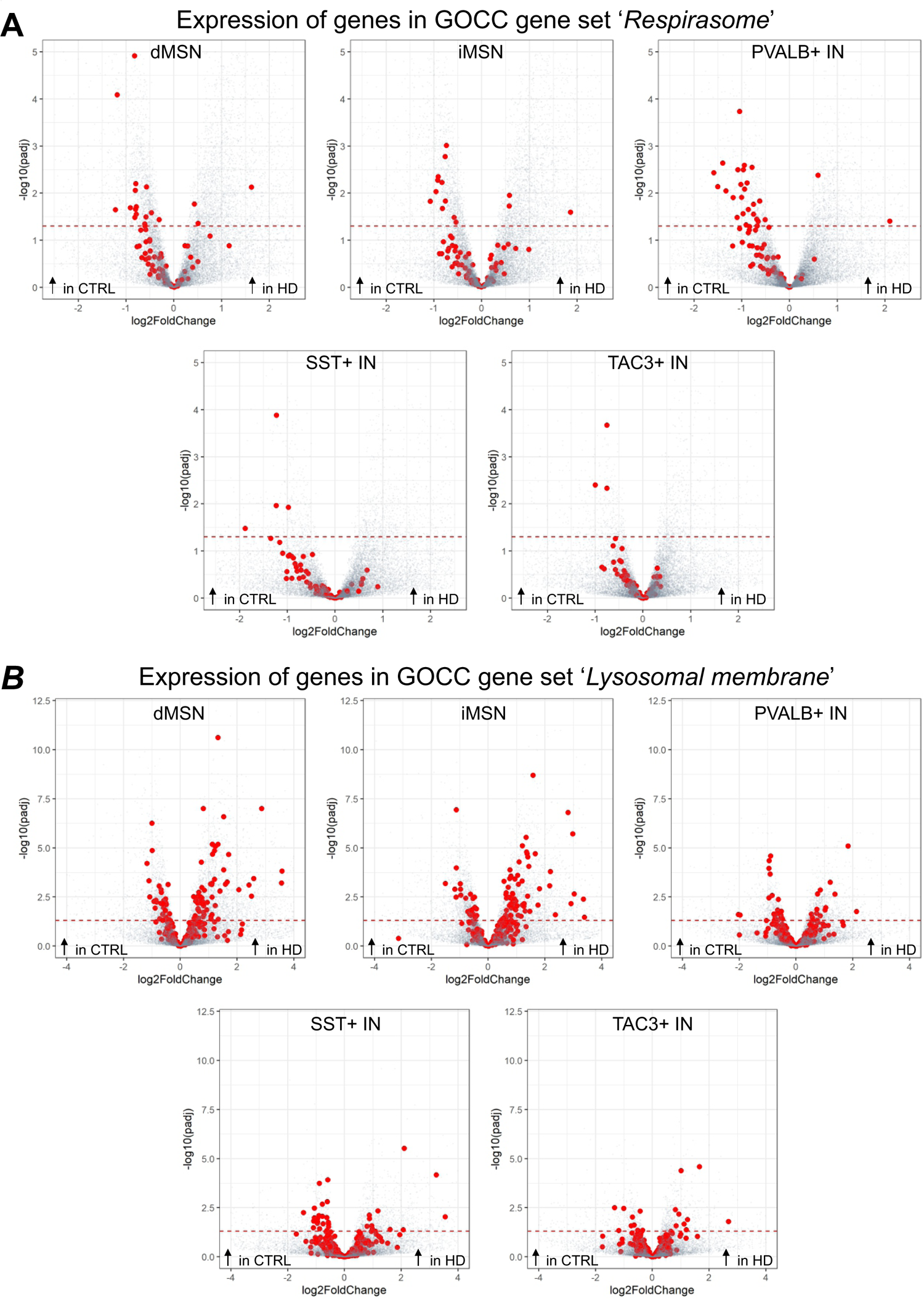
Disease-associated changes in genes associated with GO Cellular Component terms (A) ‘Respirasome’(GO:0070469) and (B) ‘Lysosomal membrane’ (GO:0005765). Dashed line denotes the threshold of statistical significance (p adj = 0.05).

**Figure S13.**
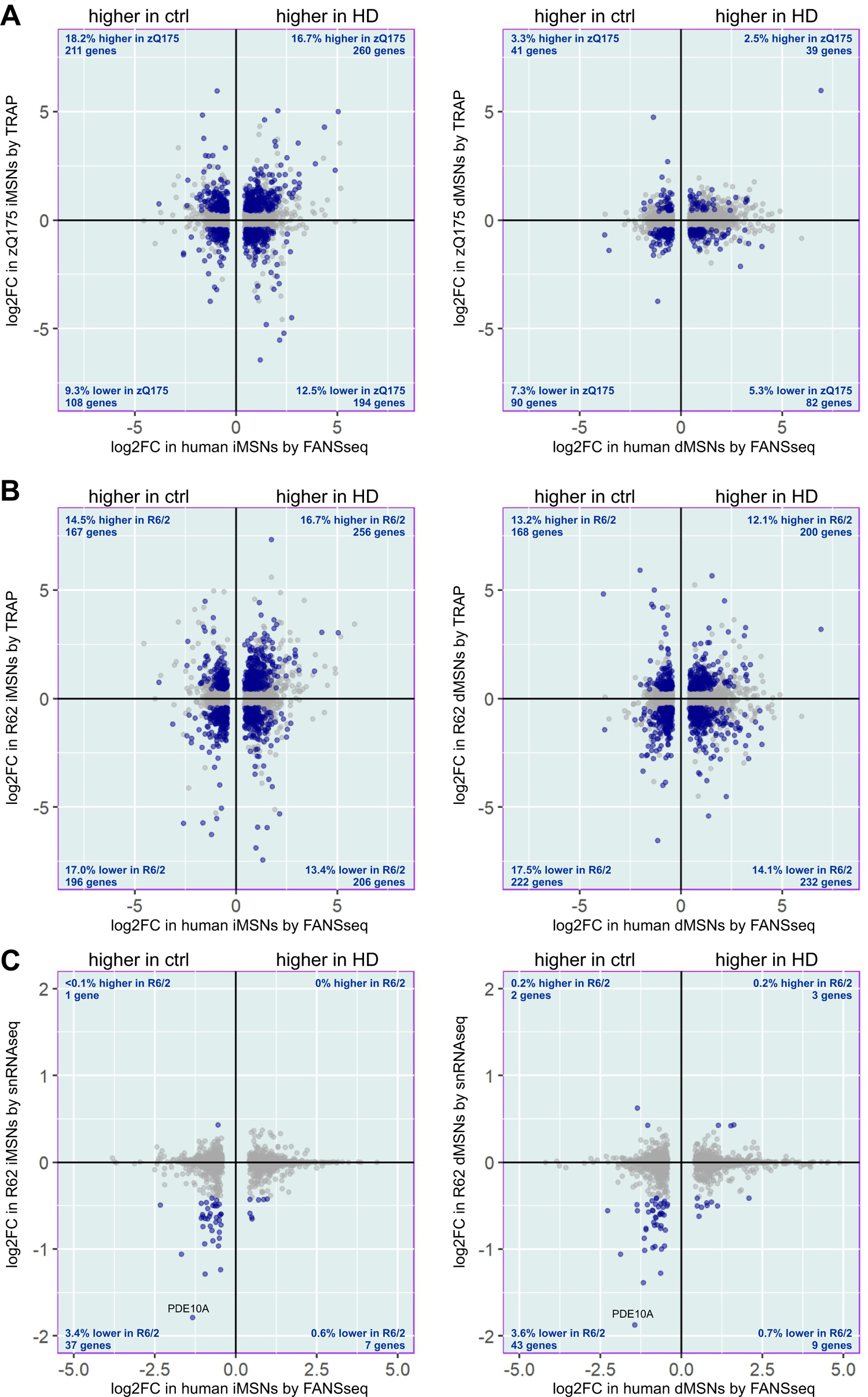
Genes with disease-associated transcript level change (p adj. < 0.05 and |log2FC|>0.415) in FANSseq data from human iMSN or dMSN were plotted against the mouse orthologue transcript level change in (**A**) TRAP data from iMSNs or dMSNs of zQ175 mice (compared to Q20 mice) and (**B**) TRAP data or (**C**) snRNAseq data from iMSNs or dMSNs of R6/2 mice (compared to wt mice)^5^. Genes with a significant transcript level change in the HD mouse model (p adj. < 0.05 and |log2FC|>0.415) are marked with blue color. Note that for most of these genes in mouse there is no significant transcript level change according to these criteria.

**Figure S14.**
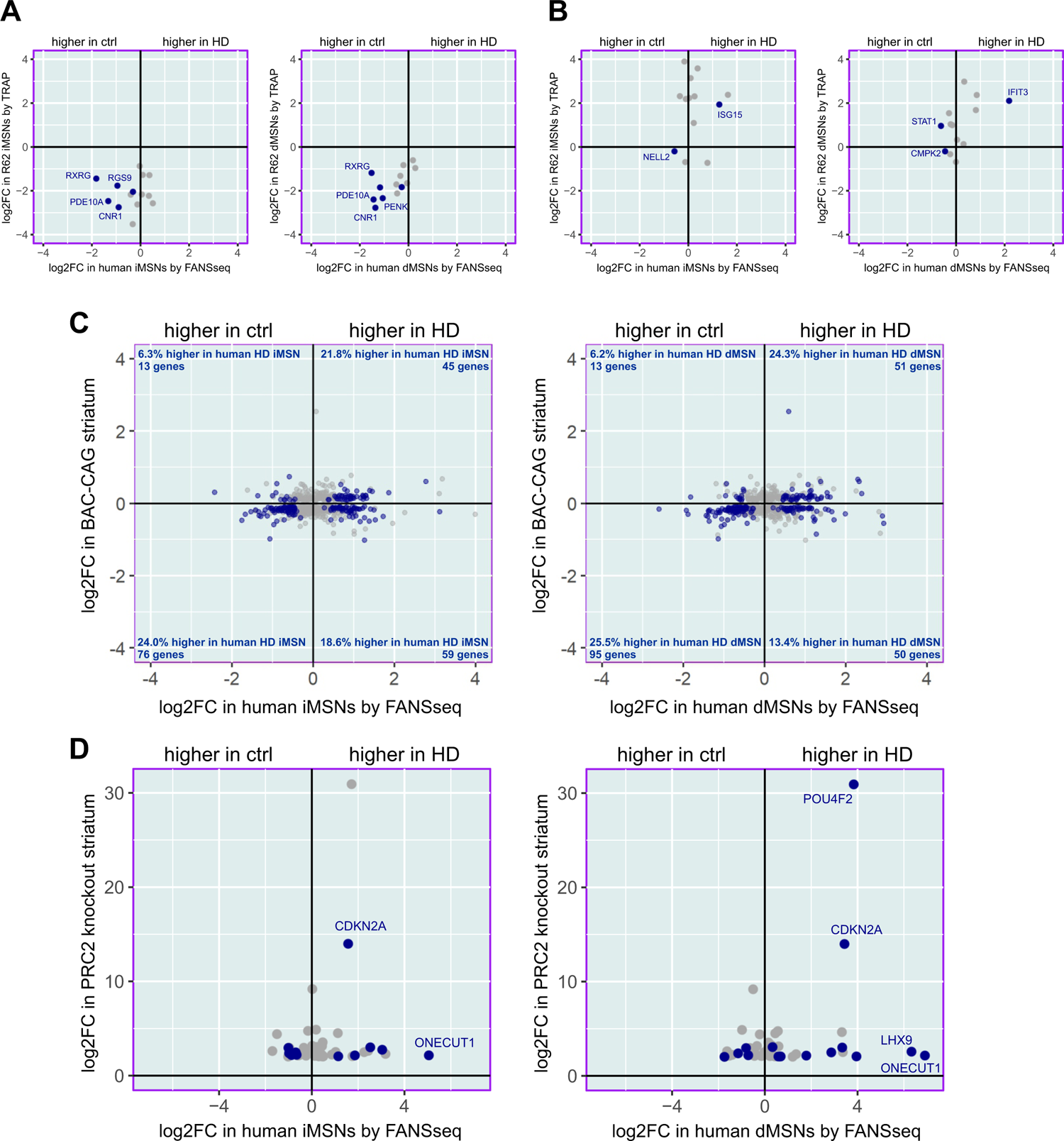
HD-associated transcript level change in human orthologues of (**A**) MSN marker genes downregulated in R6/2 mice, (**B**) innate immunity genes induced in R6/2 mice, (**C**) genes with altered expression in the striatum of BAC-CAG mice (FDR < 0.1), and (**D**) genes induced in the striatum of MSN-specific Ezh1/Ezh2 double knockout mice. The comparisons exclude genes for which none of their annotated TSS positions overlapped with ATAC-seq consensus peaks defined from control or HD donor MSN data. Genes with a significant transcript level change in human FANSseq data (p adj. < 0.05) are marked with blue color. Note that for most of these genes there is no significant transcript level change in the nuclei of human MSNs.

**Figure S15.**
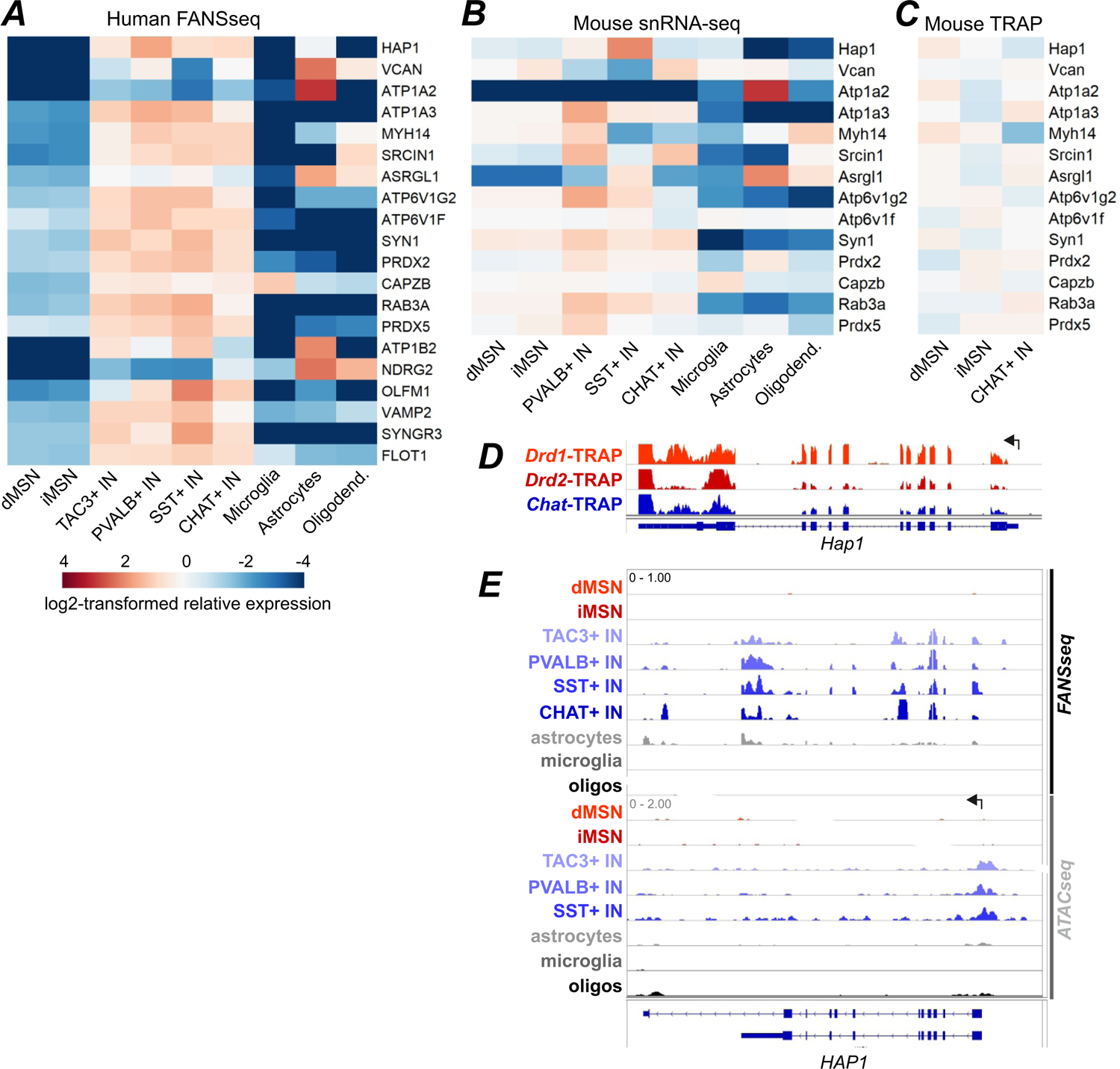
HTT interactome genes with low expression in human MSNs. We analyzed cell type-specific expression of genes identified through proteomic analysis of HTT-interacting proteins (HIPs) immunoaffinity-purified from mouse striatum^6^. (**A-C**) For many of those HIP genes that have a much lower transcript level in MSNs compared to any striatal interneuron type, this expression pattern could not have been predicted from published cell type-specific gene expression analysis of mouse striatum^3,7^. (**A**) Heatmap shows gene expression in the putamen of human control donors and includes genes for which P adj. <0.05 and log2 of fold change < −1 (by DESeq2) when comparing expression in dMSN or iMSN to any interneuron type in both putamen and caudate nucleus. (**B-C**) Genes for which lower expression in human MSNs was not predicted by (**B**) snRNA-seq data from mouse striatum^3^ or (**C**) by TRAP RNA-seq data from mouse CHAT+ INs (Chat-TRAP mouse line) and MSNs (Drd1-TRAP and Drd2-TRAP lines)^7^. N=3 for all TRAP lines. Heatmaps show log2-transformed relative expression in each cell type, calculated based on DESeq2-normalized counts (for **A** and **C**) or ‘transcripts per 100,000’ values (for **B**). (**D-E**) The gene of Huntingtin-interacting protein HAP1^8^ is transcribed in mouse MSNs and depletion of HAP1 leads to loss of MSNs in a mouse model of HD^9^. (**D**) Distribution of TRAP-RNAseq reads mapped to *Hap1*, showing its expression in MSNs and CHAT+ INs of mouse striatum. However, in humans, *HAP1* transcripts are virtually absent in MSNs, yet clearly present in striatal interneurons. These observations are supported by ATACseq data indicating that *HAP1* promoter region is not accessible in human MSNs. (**E**) Distribution of human FANSseq and ATAC-seq reads mapped to *HAP1*. Arrows mark the position of annotated transcriptional start sites. The data are from a 41-year-old male control donor.

## REFERENCES

1. Gusella, J.F., Lee, J.M., and MacDonald, M.E. (2021). Huntington’s disease: nearly four decades of human molecular genetics. Hum Mol Genet 30, R254–R263. 10.1093/hmg/ddab170.

2. Ellis, N., Tee, A., McAllister, B., Massey, T., McLauchlan, D., Stone, T., Correia, K., Loupe, J., Kim, K.H., Barker, D., et al. (2020). Genetic Risk Underlying Psychiatric and Cognitive Symptoms in Huntington’s Disease. Biol Psychiatry 87, 857–865. 10.1016/j.biopsych.2019.12.010.

3. Genetic Modifiers of Huntington’s Disease, C. (2015). Identification of Genetic Factors that Modify Clinical Onset of Huntington’s Disease. Cell 162, 516–526. 10.1016/j.cell.2015.07.003.

4. Heinsen, H., Strik, M., Bauer, M., Luther, K., Ulmar, G., Gangnus, D., Jungkunz, G., Eisenmenger, W., and Gotz, M. (1994). Cortical and striatal neurone number in Huntington’s disease. Acta Neuropathol 88, 320–333. 10.1007/BF00310376.

5. Ferrante, R.J., Kowall, N.W., Beal, M.F., Martin, J.B., Bird, E.D., and Richardson, E.P., Jr. (1987). Morphologic and histochemical characteristics of a spared subset of striatal neurons in Huntington’s disease. J Neuropathol Exp Neurol 46, 12–27. 10.1097/00005072-198701000-00002.

6. Massouh, M., Wallman, M.J., Pourcher, E., and Parent, A. (2008). The fate of the large striatal interneurons expressing calretinin in Huntington’s disease. Neurosci Res 62, 216–224. 10.1016/j.neures.2008.08.007.

7. Cicchetti, F., Prensa, L., Wu, Y., and Parent, A. (2000). Chemical anatomy of striatal interneurons in normal individuals and in patients with Huntington’s disease. Brain Res Brain Res Rev 34, 80–101. 10.1016/s0165-0173(00)00039-4.

8. DiFiglia, M., Sapp, E., Chase, K.O., Davies, S.W., Bates, G.P., Vonsattel, J.P., and Aronin, N. (1997). Aggregation of huntingtin in neuronal intranuclear inclusions and dystrophic neurites in brain. Science 277, 1990–1993. 10.1126/science.277.5334.1990.

9. Landwehrmeyer, G.B., McNeil, S.M., Dure, L.S.t., Ge, P., Aizawa, H., Huang, Q., Ambrose, C.M., Duyao, M.P., Bird, E.D., Bonilla, E., and, et al. (1995). Huntington’s disease gene: regional and cellular expression in brain of normal and affected individuals. Ann Neurol 37, 218–230. 10.1002/ana.410370213.

10. A novel gene containing a trinucleotide repeat that is expanded and unstable on Huntington’s disease chromosomes. The Huntington’s Disease Collaborative Research Group. (1993). Cell 72, 971–983. 10.1016/0092-8674(93)90585-e.

11. Telenius, H., Kremer, B., Goldberg, Y.P., Theilmann, J., Andrew, S.E., Zeisler, J., Adam, S., Greenberg, C., Ives, E.J., Clarke, L.A., and, et al. (1994). Somatic and gonadal mosaicism of the Huntington disease gene CAG repeat in brain and sperm. Nat Genet 6, 409–414. 10.1038/ng0494-409.

12. Kennedy, L., Evans, E., Chen, C.M., Craven, L., Detloff, P.J., Ennis, M., and Shelbourne, P.F. (2003). Dramatic tissue-specific mutation length increases are an early molecular event in Huntington disease pathogenesis. Hum Mol Genet 12, 3359–3367. 10.1093/hmg/ddg352.

13. Genetic Modifiers of Huntington’s Disease Consortium. Electronic address, g.h.m.h.e., and Genetic Modifiers of Huntington’s Disease, C. (2019). CAG Repeat Not Polyglutamine Length Determines Timing of Huntington’s Disease Onset. Cell 178, 887–900 e814. 10.1016/j.cell.2019.06.036.

14. Pinto, R.M., Dragileva, E., Kirby, A., Lloret, A., Lopez, E., St Claire, J., Panigrahi, G.B., Hou, C., Holloway, K., Gillis, T., et al. (2013). Mismatch repair genes Mlh1 and Mlh3 modify CAG instability in Huntington’s disease mice: genome-wide and candidate approaches. PLoS Genet 9, e1003930. 10.1371/journal.pgen.1003930.

15. Dragileva, E., Hendricks, A., Teed, A., Gillis, T., Lopez, E.T., Friedberg, E.C., Kucherlapati, R., Edelmann, W., Lunetta, K.L., MacDonald, M.E., and Wheeler, V.C. (2009). Intergenerational and striatal CAG repeat instability in Huntington’s disease knock-in mice involve different DNA repair genes. Neurobiol Dis 33, 37–47. 10.1016/j.nbd.2008.09.014.

16. Wheeler, V.C., Lebel, L.A., Vrbanac, V., Teed, A., te Riele, H., and MacDonald, M.E. (2003). Mismatch repair gene Msh2 modifies the timing of early disease in Hdh(Q111) striatum. Hum Mol Genet 12, 273–281. 10.1093/hmg/ddg056.

17. Hong, E.P., MacDonald, M.E., Wheeler, V.C., Jones, L., Holmans, P., Orth, M., Monckton, D.G., Long, J.D., Kwak, S., Gusella, J.F., and Lee, J.M. (2021). Huntington’s Disease Pathogenesis: Two Sequential Components. J Huntingtons Dis 10, 35–51. 10.3233/JHD-200427.

18. Xu, X., Stoyanova, E.I., Lemiesz, A.E., Xing, J., Mash, D.C., and Heintz, N. (2018). Species and cell-type properties of classically defined human and rodent neurons and glia. Elife 7. 10.7554/eLife.37551.

19. Krienen, F.M., Goldman, M., Zhang, Q., R, C.H.D.R., Florio, M., Machold, R., Saunders, A., Levandowski, K., Zaniewski, H., Schuman, B., et al. (2020). Innovations present in the primate interneuron repertoire. Nature 586, 262–269. 10.1038/s41586-020-2781-z.

20. Doyle, J.P., Dougherty, J.D., Heiman, M., Schmidt, E.F., Stevens, T.R., Ma, G., Bupp, S., Shrestha, P., Shah, R.D., Doughty, M.L., et al. (2008). Application of a translational profiling approach for the comparative analysis of CNS cell types. Cell 135, 749–762. 10.1016/j.cell.2008.10.029.

21. Caglayan, E., Liu, Y., and Konopka, G. (2022). Neuronal ambient RNA contamination causes misinterpreted and masked cell types in brain single-nuclei datasets. Neuron 110, 4043–4056 e4045. 10.1016/j.neuron.2022.09.010.

22. Buenrostro, J.D., Giresi, P.G., Zaba, L.C., Chang, H.Y., and Greenleaf, W.J. (2013). Transposition of native chromatin for fast and sensitive epigenomic profiling of open chromatin, DNA-binding proteins and nucleosome position. Nat Methods 10, 1213–1218. 10.1038/nmeth.2688.

23. Thornton, C.A., Johnson, K., and Moxley, R.T 3rd., (1994). Myotonic dystrophy patients have larger CTG expansions in skeletal muscle than in leukocytes. Ann Neurol 35, 104–107. 10.1002/ana.410350116.

24. Long, A., Napierala, J.S., Polak, U., Hauser, L., Koeppen, A.H., Lynch, D.R., and Napierala, M. (2017). Somatic instability of the expanded GAA repeats in Friedreich’s ataxia. PLoS One 12, e0189990. 10.1371/journal.pone.0189990.

25. Mouro Pinto, R., Arning, L., Giordano, J.V., Razghandi, P., Andrew, M.A., Gillis, T., Correia, K., Mysore, J.S., Grote Urtubey, D.M., Parwez, C.R., et al. (2020). Patterns of CAG repeat instability in the central nervous system and periphery in Huntington’s disease and in spinocerebellar ataxia type 1. Hum Mol Genet 29, 2551–2567. 10.1093/hmg/ddaa139.

26. Lokanga, R.A., Entezam, A., Kumari, D., Yudkin, D., Qin, M., Smith, C.B., and Usdin, K. (2013). Somatic expansion in mouse and human carriers of fragile X premutation alleles. Hum Mutat 34, 157–166. 10.1002/humu.22177.

27. Shelbourne, P.F., Keller-McGandy, C., Bi, W.L., Yoon, S.R., Dubeau, L., Veitch, N.J., Vonsattel, J.P., Wexler, N.S., Group, U.S.-V.C.R., Arnheim, N., and Augood, S.J. (2007). Triplet repeat mutation length gains correlate with cell-type specific vulnerability in Huntington disease brain. Hum Mol Genet 16, 1133–1142. 10.1093/hmg/ddm054.

28. Ciosi, M., Maxwell, A., Cumming, S.A., Hensman Moss, D.J., Alshammari, A.M., Flower, M.D., Durr, A., Leavitt, B.R., Roos, R.A.C., team, T.-H., et al. (2019). A genetic association study of glutamine-encoding DNA sequence structures, somatic CAG expansion, and DNA repair gene variants, with Huntington disease clinical outcomes. EBioMedicine 48, 568–580. 10.1016/j.ebiom.2019.09.020.

29. Reiner, A., Albin, R.L., Anderson, K.D., D’Amato, C.J., Penney, J.B., and Young, A.B. (1988). Differential loss of striatal projection neurons in Huntington disease. Proc Natl Acad Sci U S A 85, 5733–5737. 10.1073/pnas.85.15.5733.

30. Paulson, H. (2018). Repeat expansion diseases. Handb Clin Neurol 147, 105–123. 10.1016/B978-0-444-63233-3.00009-9.

31. Franklin, G.L., Camargo, C.H.F., Meira, A.T., Pavanelli, G.M., Milano, S.S., Germiniani, F.B., Lima, N.S.C., Raskin, S., Barsottini, O.G.P., Pedroso, J.L., et al. (2020). Is Ataxia an Underestimated Symptom of Huntington’s Disease? Front Neurol 11, 571843. 10.3389/fneur.2020.571843.

32. Singh-Bains, M.K., Mehrabi, N.F., Sehji, T., Austria, M.D.R., Tan, A.Y.S., Tippett, L.J., Dragunow, M., Waldvogel, H.J., and Faull, R.L.M. (2019). Cerebellar degeneration correlates with motor symptoms in Huntington disease. Ann Neurol 85, 396–405. 10.1002/ana.25413.

33. Nakamori, M., Pearson, C.E., and Thornton, C.A. (2011). Bidirectional transcription stimulates expansion and contraction of expanded (CTG)*(CAG) repeats. Hum Mol Genet 20, 580–588. 10.1093/hmg/ddq501.

34. Wheeler, V.C., and Dion, V. (2021). Modifiers of CAG/CTG Repeat Instability: Insights from Mammalian Models. J Huntingtons Dis 10, 123–148. 10.3233/JHD-200426.

35. Flower, M., Lomeikaite, V., Ciosi, M., Cumming, S., Morales, F., Lo, K., Hensman Moss, D., Jones, L., Holmans, P., Investigators, T.-H., et al. (2019). MSH3 modifies somatic instability and disease severity in Huntington’s and myotonic dystrophy type 1. Brain 142, 1876–1886. 10.1093/brain/awz115.

36. Manley, K., Shirley, T.L., Flaherty, L., and Messer, A. (1999). Msh2 deficiency prevents in vivo somatic instability of the CAG repeat in Huntington disease transgenic mice. Nat Genet 23, 471–473. 10.1038/70598.

37. Kovalenko, M., Dragileva, E., St Claire, J., Gillis, T., Guide, J.R., New, J., Dong, H., Kucherlapati, R., Kucherlapati, M.H., Ehrlich, M.E., et al. (2012). Msh2 acts in medium-spiny striatal neurons as an enhancer of CAG instability and mutant huntingtin phenotypes in Huntington’s disease knock-in mice. PLoS One 7, e44273. 10.1371/journal.pone.0044273.

38. Tome, S., Manley, K., Simard, J.P., Clark, G.W., Slean, M.M., Swami, M., Shelbourne, P.F., Tillier, E.R., Monckton, D.G., Messer, A., and Pearson, C.E. (2013). MSH3 polymorphisms and protein levels affect CAG repeat instability in Huntington’s disease mice. PLoS Genet 9, e1003280. 10.1371/journal.pgen.1003280.

39. Kim, K.H., Hong, E.P., Shin, J.W., Chao, M.J., Loupe, J., Gillis, T., Mysore, J.S., Holmans, P., Jones, L., Orth, M., et al. (2020). Genetic and Functional Analyses Point to FAN1 as the Source of Multiple Huntington Disease Modifier Effects. Am J Hum Genet 107, 96–110. 10.1016/j.ajhg.2020.05.012.

40. Deshmukh, A.L., Porro, A., Mohiuddin, M., Lanni, S., Panigrahi, G.B., Caron, M.C., Masson, J.Y., Sartori, A.A., and Pearson, C.E. (2021). FAN1, a DNA Repair Nuclease, as a Modifier of Repeat Expansion Disorders. J Huntingtons Dis 10, 95–122. 10.3233/JHD-200448.

41. Loupe, J.M., Pinto, R.M., Kim, K.H., Gillis, T., Mysore, J.S., Andrew, M.A., Kovalenko, M., Murtha, R., Seong, I., Gusella, J.F., et al. (2020). Promotion of somatic CAG repeat expansion by Fan1 knock-out in Huntington’s disease knock-in mice is blocked by Mlh1 knock-out. Hum Mol Genet 29, 3044–3053. 10.1093/hmg/ddaa196.

42. Goold, R., Flower, M., Moss, D.H., Medway, C., Wood-Kaczmar, A., Andre, R., Farshim, P., Bates, G.P., Holmans, P., Jones, L., and Tabrizi, S.J. (2019). FAN1 modifies Huntington’s disease progression by stabilizing the expanded HTT CAG repeat. Hum Mol Genet 28, 650–661. 10.1093/hmg/ddy375.

43. McAllister, B., Donaldson, J., Binda, C.S., Powell, S., Chughtai, U., Edwards, G., Stone, J., Lobanov, S., Elliston, L., Schuhmacher, L.N., et al. (2022). Exome sequencing of individuals with Huntington’s disease implicates FAN1 nuclease activity in slowing CAG expansion and disease onset. Nat Neurosci 25, 446–457. 10.1038/s41593-022-01033-5.

44. Deshmukh, A.L., Caron, M.C., Mohiuddin, M., Lanni, S., Panigrahi, G.B., Khan, M., Engchuan, W., Shum, N., Faruqui, A., Wang, P., et al. (2021). FAN1 exo-not endo-nuclease pausing on disease-associated slipped-DNA repeats: A mechanism of repeat instability. Cell Rep 37, 110078. 10.1016/j.celrep.2021.110078.

45. Lee, H., Fenster, R.J., Pineda, S.S., Gibbs, W.S., Mohammadi, S., Davila-Velderrain, J., Garcia, F.J., Therrien, M., Novis, H.S., Gao, F., et al. (2020). Cell Type-Specific Transcriptomics Reveals that Mutant Huntingtin Leads to Mitochondrial RNA Release and Neuronal Innate Immune Activation. Neuron 107, 891–908 e898. 10.1016/j.neuron.2020.06.021.

46. Sardiello, M., Palmieri, M., di Ronza, A., Medina, D.L., Valenza, M., Gennarino, V.A., Di Malta, C., Donaudy, F., Embrione, V., Polishchuk, R.S., et al. (2009). A gene network regulating lysosomal biogenesis and function. Science 325, 473–477. 10.1126/science.1174447.

47. Rui, Y.N., Xu, Z., Patel, B., Chen, Z., Chen, D., Tito, A., David, G., Sun, Y., Stimming, E.F., Bellen, H.J., et al. (2015). Huntingtin functions as a scaffold for selective macroautophagy. Nat Cell Biol 17, 262–275. 10.1038/ncb3101.

48. Cason, S.E., Carman, P.J., Van Duyne, C., Goldsmith, J., Dominguez, R., and Holzbaur, E.L.F. (2021). Sequential dynein effectors regulate axonal autophagosome motility in a maturation-dependent pathway. J Cell Biol 220. 10.1083/jcb.202010179.

49. Lee, J.M., Huang, Y., Orth, M., Gillis, T., Siciliano, J., Hong, E., Mysore, J.S., Lucente, D., Wheeler, V.C., Seong, I.S., et al. (2022). Genetic modifiers of Huntington disease differentially influence motor and cognitive domains. Am J Hum Genet 109, 885–899. 10.1016/j.ajhg.2022.03.004.

50. Christmann, M., and Kaina, B. (2013). Transcriptional regulation of human DNA repair genes following genotoxic stress: trigger mechanisms, inducible responses and genotoxic adaptation. Nucleic Acids Res 41, 8403–8420. 10.1093/nar/gkt635.

51. Wertz, M.H., Mitchem, M.R., Pineda, S.S., Hachigian, L.J., Lee, H., Lau, V., Powers, A., Kulicke, R., Madan, G.K., Colic, M., et al. (2020). Genome-wide In Vivo CNS Screening Identifies Genes that Modify CNS Neuronal Survival and mHTT Toxicity. Neuron 106, 76–89 e78. 10.1016/j.neuron.2020.01.004.

52. Ito, N., Hendriks, W.T., Dhakal, J., Vaine, C.A., Liu, C., Shin, D., Shin, K., Wakabayashi-Ito, N., Dy, M., Multhaupt-Buell, T., et al. (2016). Decreased N-TAF1 expression in X-linked dystonia-parkinsonism patient-specific neural stem cells. Dis Model Mech 9, 451–462. 10.1242/dmm.022590.

53. Makino, S., Kaji, R., Ando, S., Tomizawa, M., Yasuno, K., Goto, S., Matsumoto, S., Tabuena, M.D., Maranon, E., Dantes, M., et al. (2007). Reduced neuron-specific expression of the TAF1 gene is associated with X-linked dystonia-parkinsonism. Am J Hum Genet 80, 393–406. 10.1086/512129.

54. Campion, L.N., Mejia Maza, A., Yadav, R., Penney, E.B., Murcar, M.G., Correia, K., Gillis, T., Fernandez-Cerado, C., Velasco-Andrada, M.S., Legarda, G.P., et al. (2022). Tissue-specific and repeat length-dependent somatic instability of the X-linked dystonia parkinsonism-associated CCCTCT repeat. Acta Neuropathol Commun 10, 49. 10.1186/s40478-022-01349-0.

55. Veldman, M.B., and Yang, X.W. (2018). Molecular insights into cortico-striatal miscommunications in Huntington’s disease. Curr Opin Neurobiol 48, 79–89. 10.1016/j.conb.2017.10.019.

56. Gu, X., Richman, J., Langfelder, P., Wang, N., Zhang, S., Banez-Coronel, M., Wang, H.B., Yang, L., Ramanathan, L., Deng, L., et al. (2022). Uninterrupted CAG repeat drives striatum-selective transcriptionopathy and nuclear pathogenesis in human Huntingtin BAC mice. Neuron 110, 1173–1192 e1177. 10.1016/j.neuron.2022.01.006.

57. von Schimmelmann, M., Feinberg, P.A., Sullivan, J.M., Ku, S.M., Badimon, A., Duff, M.K., Wang, Z., Lachmann, A., Dewell, S., Ma’ayan, A., et al. (2016). Polycomb repressive complex 2 (PRC2) silences genes responsible for neurodegeneration. Nat Neurosci 19, 1321–1330. 10.1038/nn.4360.

58. Suzuki, M., Desmond, T.J., Albin, R.L., and Frey, K.A. (2001). Vesicular neurotransmitter transporters in Huntington’s disease: initial observations and comparison with traditional synaptic markers. Synapse 41, 329–336. 10.1002/syn.1089.

59. Rub, U., Hoche, F., Brunt, E.R., Heinsen, H., Seidel, K., Del Turco, D., Paulson, H.L., Bohl, J., von Gall, C., Vonsattel, J.P., et al. (2013). Degeneration of the cerebellum in Huntington’s disease (HD): possible relevance for the clinical picture and potential gateway to pathological mechanisms of the disease process. Brain Pathol 23, 165–177. 10.1111/j.1750-3639.2012.00629.x.

60. Jeste, D.V., Barban, L., and Parisi, J. (1984). Reduced Purkinje cell density in Huntington’s disease. Exp Neurol 85, 78–86. 10.1016/0014-4886(84)90162-6.

61. Tian, L., Gu, L., and Li, G.M. (2009). Distinct nucleotide binding/hydrolysis properties and molar ratio of MutSalpha and MutSbeta determine their differential mismatch binding activities. J Biol Chem 284, 11557–11562. 10.1074/jbc.M900908200.

62. Brown, M.W., Kim, Y., Williams, G.M., Huck, J.D., Surtees, J.A., and Finkelstein, I.J. (2016). Dynamic DNA binding licenses a repair factor to bypass roadblocks in search of DNA lesions. Nat Commun 7, 10607. 10.1038/ncomms10607.

63. Britton, B.M., London, J.A., Martin-Lopez, J., Jones, N.D., Liu, J., Lee, J.B., and Fishel, R. (2022). Exploiting the distinctive properties of the bacterial and human MutS homolog sliding clamps on mismatched DNA. J Biol Chem 298, 102505. 10.1016/j.jbc.2022.102505.

64. Gupta, S., Gellert, M., and Yang, W. (2011). Mechanism of mismatch recognition revealed by human MutSbeta bound to unpaired DNA loops. Nat Struct Mol Biol 19, 72–78. 10.1038/nsmb.2175.

65. Owen, B.A., Yang, Z., Lai, M., Gajec, M., Badger, J.D, 2nd., Hayes, J.J., Edelmann, W., Kucherlapati, R., Wilson, T.M., and McMurray, C.T. (2005). (CAG)(n)-hairpin DNA binds to Msh2-Msh3 and changes properties of mismatch recognition. Nat Struct Mol Biol 12, 663-670. 10.1038/nsmb965.

66. Koeppen, A.H. (2018). The Neuropathology of Spinocerebellar Ataxia Type 3/Machado-Joseph Disease. Adv Exp Med Biol 1049, 233–241. 10.1007/978-3-319-71779-1_11.

67. Rousseaux, M.W.C., Tschumperlin, T., Lu, H.C., Lackey, E.P., Bondar, V.V., Wan, Y.W., Tan, Q., Adamski, C.J., Friedrich, J., Twaroski, K., et al. (2018). ATXN1-CIC Complex Is the Primary Driver of Cerebellar Pathology in Spinocerebellar Ataxia Type 1 through a Gain-of-Function Mechanism. Neuron 97, 1235–1243 e1235. 10.1016/j.neuron.2018.02.013.

68. Greco, T.M., Secker, C., Ramos, E.S., Federspiel, J.D., Liu, J.P., Perez, A.M., Al-Ramahi, I., Cantle, J.P., Carroll, J.B., Botas, J., et al. (2022). Dynamics of huntingtin protein interactions in the striatum identifies candidate modifiers of Huntington disease. Cell Syst 13, 304–320 e305. 10.1016/j.cels.2022.01.005.

69. Brandebura, A.N., Paumier, A., Onur, T.S., and Allen, N.J. (2023). Astrocyte contribution to dysfunction, risk and progression in neurodegenerative disorders. Nat Rev Neurosci 24, 23–39. 10.1038/s41583-022-00641-1.

70. Hickman, S., Izzy, S., Sen, P., Morsett, L., and El Khoury, J. (2018). Microglia in neurodegeneration. Nat Neurosci 21, 1359–1369. 10.1038/s41593-018-0242-x.

71. Liao, Y., Smyth, G.K., and Shi, W. (2013). The Subread aligner: fast, accurate and scalable read mapping by seed-and-vote. Nucleic Acids Res 41, e108. 10.1093/nar/gkt214.

72. Zhang, Y., Liu, T., Meyer, C.A., Eeckhoute, J., Johnson, D.S., Bernstein, B.E., Nusbaum, C., Myers, R.M., Brown, M., Li, W., and Liu, X.S. (2008). Model-based analysis of ChIP-Seq (MACS). Genome Biol 9, R137. 10.1186/gb-2008-9-9-r137.

73. Wang, Q., Li, M., Wu, T., Zhan, L., Li, L., Chen, M., Xie, W., Xie, Z., Hu, E., Xu, S., and Yu, G. (2022). Exploring Epigenomic Datasets by ChIPseeker. Curr Protoc 2, e585. 10.1002/cpz1.585.

74. Marini, F., and Binder, H. (2019). pcaExplorer: an R/Bioconductor package for interacting with RNA-seq principal components. BMC Bioinformatics 20, 331. 10.1186/s12859-019-2879-1.

75. Love, M.I., Hogenesch, J.B., and Irizarry, R.A. (2016). Modeling of RNA-seq fragment sequence bias reduces systematic errors in transcript abundance estimation. Nat Biotechnol 34, 1287–1291. 10.1038/nbt.3682.

76. Robinson, J.T., Thorvaldsdottir, H., Winckler, W., Guttman, M., Lander, E.S., Getz, G., and Mesirov, J.P. (2011). Integrative genomics viewer. Nat Biotechnol 29, 24–26. 10.1038/nbt.1754.

77. Yu, G., Wang, L.G., Han, Y., and He, Q.Y. (2012). clusterProfiler: an R package for comparing biological themes among gene clusters. OMICS 16, 284–287. 10.1089/omi.2011.0118.

78. Warner, J.H., Long, J.D., Mills, J.A., Langbehn, D.R., Ware, J., Mohan, A., and Sampaio, C. (2022). Standardizing the CAP Score in Huntington’s Disease by Predicting Age-at-Onset. J Huntingtons Dis 11, 153–171. 10.3233/JHD-210475.

79. Feng, J., Liu, T., Qin, B., Zhang, Y., and Liu, X.S. (2012). Identifying ChIP-seq enrichment using MACS. Nat Protoc 7, 1728–1740. 10.1038/nprot.2012.101.

80. Love, M.I., Soneson, C., and Patro, R. (2018). Swimming downstream: statistical analysis of differential transcript usage following Salmon quantification. F1000Res 7, 952. 10.12688/f1000research.15398.3.

81. Maity, R., Pauty, J., Krietsch, J., Buisson, R., Genois, M.M., and Masson, J.Y. (2013). GST-His purification: a two-step affinity purification protocol yielding full-length purified proteins. J Vis Exp, e50320. 10.3791/50320.

82. Pearson, C.E., Ewel, A., Acharya, S., Fishel, R.A., and Sinden, R.R. (1997). Human MSH2 binds to trinucleotide repeat DNA structures associated with neurodegenerative diseases. Hum Mol Genet 6, 1117–1123. 10.1093/hmg/6.7.1117.

83. Panigrahi, G.B., Slean, M.M., Simard, J.P., Gileadi, O., and Pearson, C.E. (2010). Isolated short CTG/CAG DNA slip-outs are repaired efficiently by hMutSbeta, but clustered slip-outs are poorly repaired. Proc Natl Acad Sci U S A 107, 12593–12598. 10.1073/pnas.0909087107.

84. Panigrahi, G.B., Lau, R., Montgomery, S.E., Leonard, M.R., and Pearson, C.E. (2005). Slipped (CTG)*(CAG) repeats can be correctly repaired, escape repair or undergo error-prone repair. Nat Struct Mol Biol 12, 654–662. 10.1038/nsmb959.

## SUPPLEMENTAL REFERENCES

1. Krienen, F.M., Goldman, M., Zhang, Q., R, C.H.D.R., Florio, M., Machold, R., Saunders, A., Levandowski, K., Zaniewski, H., Schuman, B., et al. (2020). Innovations present in the primate interneuron repertoire. Nature 586, 262–269. 10.1038/s41586-020-2781-z.

2. Deshmukh, A.L., Caron, M.C., Mohiuddin, M., Lanni, S., Panigrahi, G.B., Khan, M., Engchuan, W., Shum, N., Faruqui, A., Wang, P., et al. (2021). FAN1 exo-not endo-nuclease pausing on disease-associated slipped-DNA repeats: A mechanism of repeat instability. Cell Rep 37, 110078. 10.1016/j.celrep.2021.110078.

3. Saunders, A., Macosko, E.Z., Wysoker, A., Goldman, M., Krienen, F.M., de Rivera, H., Bien, E., Baum, M., Bortolin, L., Wang, S., et al. (2018). Molecular Diversity and Specializations among the Cells of the Adult Mouse Brain. Cell 174, 1015–1030 e1016. 10.1016/j.cell.2018.07.028.

4. Wertz, M.H., Mitchem, M.R., Pineda, S.S., Hachigian, L.J., Lee, H., Lau, V., Powers, A., Kulicke, R., Madan, G.K., Colic, M., et al. (2020). Genome-wide In Vivo CNS Screening Identifies Genes that Modify CNS Neuronal Survival and mHTT Toxicity. Neuron 106, 76–89 e78. 10.1016/j.neuron.2020.01.004.

5. Lee, H., Fenster, R.J., Pineda, S.S., Gibbs, W.S., Mohammadi, S., Davila-Velderrain, J., Garcia, F.J., Therrien, M., Novis, H.S., Gao, F., et al. (2020). Cell Type-Specific Transcriptomics Reveals that Mutant Huntingtin Leads to Mitochondrial RNA Release and Neuronal Innate Immune Activation. Neuron 107, 891–908 e898. 10.1016/j.neuron.2020.06.021.

6. Greco, T.M., Secker, C., Ramos, E.S., Federspiel, J.D., Liu, J.P., Perez, A.M., Al-Ramahi, I., Cantle, J.P., Carroll, J.B., Botas, J., et al. (2022). Dynamics of huntingtin protein interactions in the striatum identifies candidate modifiers of Huntington disease. Cell Syst 13, 304–320 e305. 10.1016/j.cels.2022.01.005.

7. Doyle, J.P., Dougherty, J.D., Heiman, M., Schmidt, E.F., Stevens, T.R., Ma, G., Bupp, S., Shrestha, P., Shah, R.D., Doughty, M.L., et al. (2008). Application of a translational profiling approach for the comparative analysis of CNS cell types. Cell 135, 749–762. 10.1016/j.cell.2008.10.029.

8. Li, X.J., Li, S.H., Sharp, A.H., Nucifora, F.C., Jr., Schilling, G., Lanahan, A., Worley, P., Snyder, S.H., and Ross, C.A. (1995). A huntingtin-associated protein enriched in brain with implications for pathology. Nature 378, 398–402. 10.1038/378398a0.

9. Liu, Q., Cheng, S., Yang, H., Zhu, L., Pan, Y., Jing, L., Tang, B., Li, S., and Li, X.J. (2020). Loss of Hap1 selectively promotes striatal degeneration in Huntington disease mice. Proc Natl Acad Sci U S A 117, 20265–20273. 10.1073/pnas.2002283117.

